# *In silico* evidence for the utility of parsimonious root phenotypes for improved vegetative growth and carbon sequestration under drought

**DOI:** 10.1101/2022.08.04.502508

**Authors:** Ernst D. Schäfer, Ishan Ajmera, Etienne Farcot, Markus R. Owen, Leah R. Band, Jonathan P. Lynch

**Author notes:** Correspondence: Jonathan P. Lynch. These authors contributed equally to this work and share first authorship (Ernst D. Schäfer, Ishan Ajmera).

## Abstract

Drought is a primary constraint to crop yields and climate change is expected to increase the frequency and severity of drought stress in the future. It has been hypothesized that crops can be made more resistant to drought and better able to sequester atmospheric carbon in the soil by selecting appropriate root phenotypes. We introduce *OpenSimRoot_v2*, an upgraded version of the functional-structural plant/soil model *OpenSimRoot*, and use it to test the utility of a maize root phenotype with fewer and steeper axial roots, reduced lateral root branching density, and more aerenchyma formation (i.e. the ‘Steep, Cheap, and Deep’ (SCD) ideotype) and different combinations of underlying SCD root phene states under rainfed and drought conditions in three distinct maize growing pedoclimatic environments in the USA, Nigeria, and Mexico. In all environments where plants are subjected to drought stress the SCD ideotype as well as several intermediate phenotypes lead to greater shoot biomass after 42 days. As an additional advantage, the amount of carbon deposited below 50 cm in the soil is twice as great for the SCD phenotype as for the reference phenotype in 5 out of 6 simulated environments. We conclude that crop growth and deep soil carbon deposition can be improved by breeding maize plants with fewer axial roots, reduced lateral root branching density, and more aerenchyma formation.

## 1 INTRODUCTION

Globally, the incidence and severity of droughts under climate change have been increasing over the past few decades, causing substantial agricultural losses with cascading enviro-socioeconomic impacts Mukherjee et al. (2018); Reddy and Hodges (2000). Drought is a complex phenomenon commonly characterized by a suboptimal availability of water for a sustained period Wilhite and Glantz (1985); Van Loon (2015). In an agricultural context, drought is attributed to the reduction in water supply required by the plant due to soil water deficit, mostly in the root zone, and/or an increase in water loss via transpiration Boken et al. (2005); Salehi-Lisar and Bakhshayeshan-Agdam (2016). Soil water availability and plant adaptation to water deficit are spatiotemporally influenced by an array of pedo-climatic factors. These factors pose a major challenge in understanding drought and its impact on plants, which would have implications for improving agricultural practices and breeding efforts.

From the paradigm of the soil-plant-atmosphere continuum, the plant water status reflects the synchronized response of the plant to soil water availability and atmospheric demand Silva and Lambers (2021). Shoot water loss is driven by atmospheric demand together with leaf area, stomatal conductance, and intrinsic water uptake capacity of the plant De Swaef et al. (2022). On other hand, soil properties and root architecture largely determine the water availability to the plant. To counter water deficit, plants elicit an array of response mechanisms, which broadly involve - a) limiting water loss via stomatal adjustment and limiting shoot growth, b) increasing root foraging to enhance water uptake, and c) restricting tissue dehydration via osmotic and metabolic adjustments Bray (1997); Dodd and Ryan (2016); Salehi-Lisar and Bakhshayeshan-Agdam (2016).

One avenue to counter water loss via transpiration is to increase root water capture. Since water is often available in the deeper soil profiles, in most environments deeper rooting improves drought resistance. In this context, the ‘Steep, Cheap, and Deep’ (SCD) root ideotype was proposed over a decade ago for improving drought resistance in crops, by increasing root depth and in turn water acquisition from deep soil domains Lynch (2013). Fundamentally, the SCD ideotype can be perceived as an integrated root phenotype that aims to optimize how the internal plant resources and soil foraging activities are spatiotemporally distributed among different root classes and across the plant to improve its performance in response to edaphic stresses, including drought. With the root system adapted at morphological, anatomical, and physiological scales, the SCD ideotypes that enable root foraging of deeper soil strata are typically characterized by steeper axial root angles, reduced number of axial roots, reduced lateral branching densities, and formation of root cortical aerenchyma and other anatomical phenotypes that reduce the metabolic cost of soil exploration.

Over the years, various other combinations of root phenotypes corresponding to SCD ideotypes have been proposed that, *in silico* and/or *in vivo*, have been associated with improved crop growth under suboptimal water and nitrogen availability, including in maize Lynch (2018, 2019, 2022). Reducing the number of axial roots decreases competition among axial roots for internal plant resources and external soil resources, which allows the remaining axial roots to grow deeper, increasing water capture from deeper soil layers. This was confirmed in a field study where maize plants with fewer axial roots had increased rooting depth, leaf water content, shoot biomass, and yield under drought. Nitrogen in agricultural soils is generally a mobile resource (as nitrate) that, like water, is often more available in deep soil strata Thorup-Kristensen et al. (2020).

Reduced axial root number increased shoot biomass for maize and rice in response to suboptimal nitrogen availability Saengwilai et al. (2014); Gao and Lynch (2016); Ajmera et al. (2022). When the soil gets very dry and soil hydraulic conductivity becomes extremely low, there is significant competition for water and nitrogen among neighboring lateral roots. This means that reducing lateral root branching density is beneficial for plant performance under suboptimal water and nitrogen availability Postma et al. (2014); Zhan and Lynch (2015). Furthermore, the utility of steeper axial root angles for improved tolerance to drought and low nitrogen supply by developing a deeper root system has been confirmed in different crop species Manschadi et al. (2008); Trachsel et al. (2013); Uga et al. (2013); Lynch and Wojciechowski (2015); Schneider et al. (2022). Since one of the secondary effects of water deprivation on plants is a reduction in photosynthesis, plants suffering from water shortages will have limited photosynthetically-derived carbon. Anatomical phenotypes that reduced the metabolic cost of soil exploration should therefore improve soil exploration and water capture under drought, which is the case with root cortical aerenchyma, reduced cortical cell file number, and increased cortical cell size in maize Lynch et al. (2021).

It is noteworthy that a substantial amount of photosynthate is allocated to roots and eventually deposited into the rhizosphere. Shoot-derived carbon decays twice as fast as that derived from roots. In addition, the decomposition of plant residues decreases with soil depth. Given this, root phene states that enable greater rooting depth directly increase soil carbon sequestration thereby removing atmospheric CO_2_ and concurrently, enhancing soil organic matter and quality Kell (2011, 2012); Lynch (2015).

The SCD phenotype is composed of multiple root phene states. These phene states may interact and influence the utility of each other. The interaction among different phene state varies with environments including water regimes and pedo-climatic conditions. The potential combinations of these phenes would generate multitudes of potential phenotypes with varying utilities over different environments. Given this, simulation modeling is the only feasible way to explore the ‘fitness landscape’ of root phenotypes. The functional-structural plant/soil model *OpenSimRoot* has been useful in assessing various aspects of this root phenotype, particularly under low nitrogen, phosphorus, and potassium availability for different crop species in varying environments Dathe et al. (2016); Postma et al. (2017); Rangarajan et al. (2018); Ajmera et al. (2022). One of the main features of *OpenSimRoot* is that it simulates carbon assimilation, calculates carbon requirements for growth and maintenance of simulated tissues, and responds to insufficient carbon availability by reducing growth. Changes to the root-shoot ratio in response to environmental conditions are the emergent property of the model attributed to the simulated carbon and nutrient economies.

In this work, we introduce *OpenSimRoot_v2*, which adds new functionality to *OpenSimRoot*, enabling simulation of plant and soil responses to drought. This functionality includes the combined implementation of different bio-physio-chemical models available in the literature; namely the Farquhar-von Caemmerer-Berry (FvCB) model for C_3_ and C_4_photosynthesis, leaf gas exchange and stomatal conductance Von Caemmerer and Farquhar (1981); Leuning (1995), leaf temperature and energy balance models Von Caemmerer (2000), sun/shade model for leaf-to-canopy scaling De Pury and Farquhar (1997), a model for nitrogen-limited photosynthesis Kull and Kruijt (1998), water stress response functions, and models for simulating day-night cycles and the corresponding carbon allocations Sulpice et al. (2014). With this upgrade, the *OpenSimRoot_v2* still remains backward compatible enabling implementation of earlier OpenSimRoot versions with an appropriate selection of models and functions in the input files.

To better understand the relationship between root phenotypes and drought resistance, this *in silico* study evaluates the “Steep, Cheap, and Deep” ideotype and all combinations of the associated phene states in exemplary maize growing environments in the USA, Nigeria, and Mexico with distinct pedo-climatic conditions under rainfed and terminal drought conditions. Our objectives are:

- Ascertain if the “Steep, Cheap, and Deep” ideotype is advantageous, that is, leads to greater shoot biomass, under water deficit stress (henceforth simply ‘drought stress’).
- Determine the extent to which interactions among phene states of the “Steep, Cheap, and Deep” ideotype contribute to performance under drought.
- Quantify the effects of different root phenotypes on shoot biomass, rooting depth, estimated carbon deposition at depth, and carbon use efficiency (water uptake per unit of carbon invested in roots) in rainfed and drought conditions.

## 2 MATERIALS AND METHODS

*OpenSimRoot* is a feature-rich, open-source, functional-structural plant/soil simulation model, which explicitly simulates the geometry of roots and soil properties in three dimensions over time. In *OpenSimRoot*, the shoot is simulated through a number of state variables in a non-geometrical fashion. Being modular, *OpenSimRoot* encapsulates different submodels to represent various processes such as nutrient and water acquisition, aerenchyma formation, carbon sources and sinks, and nitrogen mineralization. Plant growth is determined by potential growth rates, constraints imposed by the availability of nutrients and carbon, and modifiers depending on the environment. These particular includes soils differing in bulk density, texture, and hydraulic properties. Using the root impedance model added to *OpenSimRoot* in Strock et al. (2022), we capture the impact of soil bulk density on root growth. Also included are models that consider the effect of soil water status on soil penetration resistance, with wetter soils being easier to penetrate. This is especially important when simulating plants in drying soil since root distribution affects the distribution of soil water and vice versa. To accurately simulate the effects of drought stress on stomatal conductance, photosynthesis, and overall plant development we introduce here *OpenSimRoot_v2*, which adds submodels for various state variables related to photosynthesis. Figure 1 provides an overview of the most important submodels which were added and updated and their most important interactions. Note that many submodels and interactions were left out of this diagram for the sake of readability.

**Figure 1.**
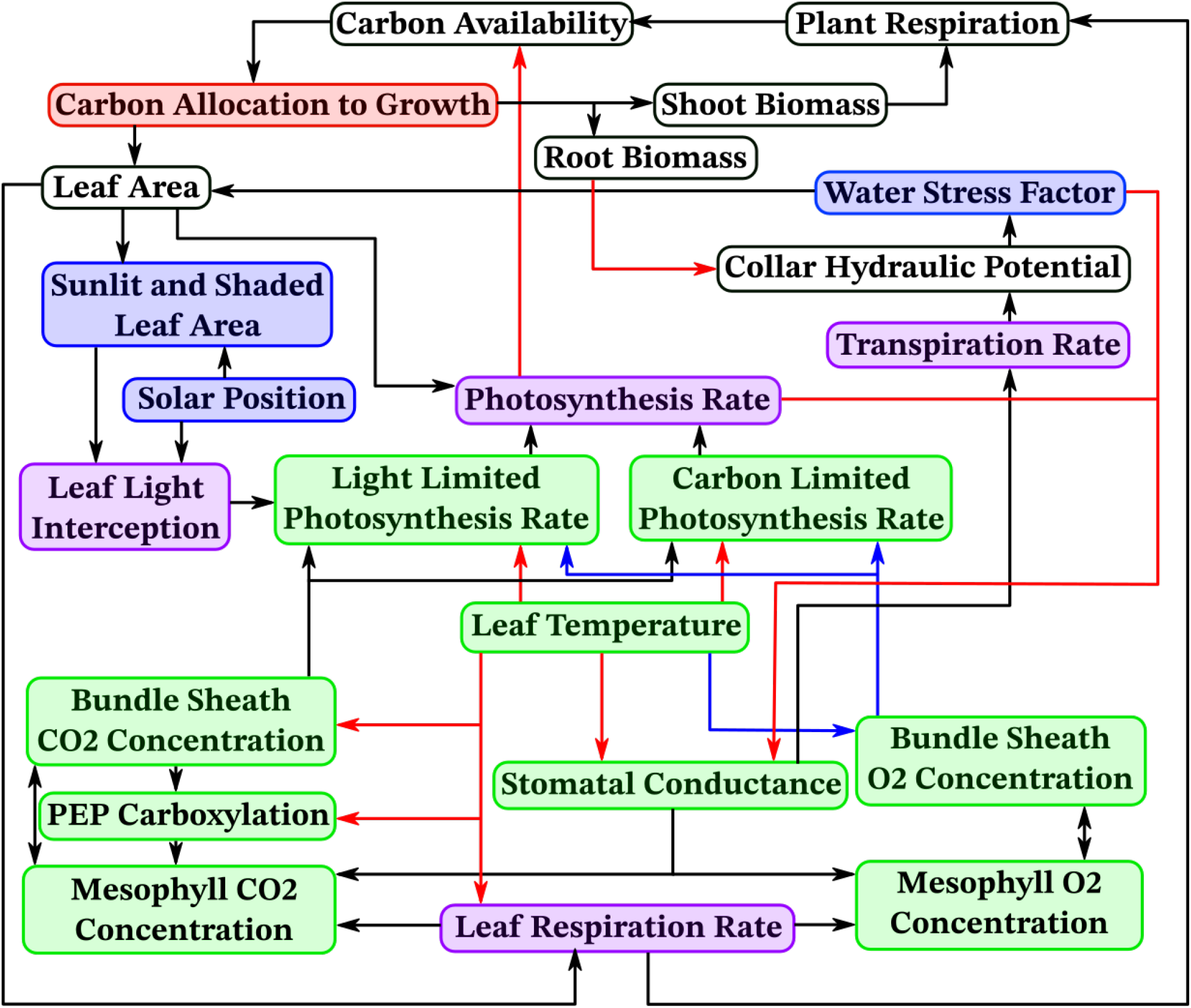
An overview of important new models and their dependencies in *OpenSimRoot*. Arrows point from a variable to variables depending on it, arrow colours are solely for distinguishing between them. White indicates models which were already present and are unchanged, red indicates models which were updated, blue indicates newly added models, green indicates newly added models with separate sunlit and shaded instances and purple indicates already present models which now have separate sunlit and shaded instances.

In *OpenSimRoot*, state variables are represented by objects which can be constant, follow a predefined variation over time, drawn from a random distribution, or calculated by a relevant class. Information moves between different parts of the model through a common application programming interface (API). Because of this modular structure, one can easily switch between different model implementations for a given state variable without the rest of the model being affected. Likewise, adding new state variables and model implementations is relatively straightforward and usually does not require any modification to the rest of the code. The new models added here required some additions to the *OpenSimRoot* engine, which we describe in section 2.9. We added new state variables and corresponding models and added updated models for some existing state variables to *OpenSimRoot*, which we describe in the following subsections. Readers are encouraged to consult the referenced papers for more details on and explanations of the model equations in these subsections.

### 2.1 Water Stress

To properly model the effect of drought on plant functioning and development we need to quantify water stress. We do this by adding a water stress minimodel similar to how nutrient stresses are captured in *OpenSimRoot*. This water stress minimodel captures water stress as a number *S*_*w*_ between 0 and 1, where 1 is no stress and 0 is maximum stress. It is known that abscisic acid (ABA) plays a role in the plant drought response Ikegami et al. (2009); Kriedemann et al. (1972); Loveys (1977); McAdam and Brodribb (2016); Okamoto et al. (2009); Sack et al. (2018), but *OpenSimRoot* does not simulate the dynamics of phytohormones neither does it explicitly capture shoot geometry to simulate leaf water potential. As a result, we use a proxy to leaf water potential to capture drought stress in *OpenSimRoot_v2*. Notably, *OpenSimRoot* does includes a xylem water flow model described in Doussan et al. (1998); Postma et al. (2017), which assumes steady-state flow to calculate the quantity and distribution of water uptake by the roots for a given hydraulic potential at the collar (which is where the hypocotyl and shoot meet). This collar hydraulic potential is set such that the water uptake by the root system matches the potential transpiration, which is either calculated using the Penman-Monteith equations Zotarelli et al. (2010) or based on photosynthesis rates. In *OpenSimRoot_v2*, we translate the collar potential Ψ_*c*_ in hPa to a stress factor *S*_*w*_(Ψ_*c*_) using a drought response curve defined in the input files. In this paper we used the following drought response curve:

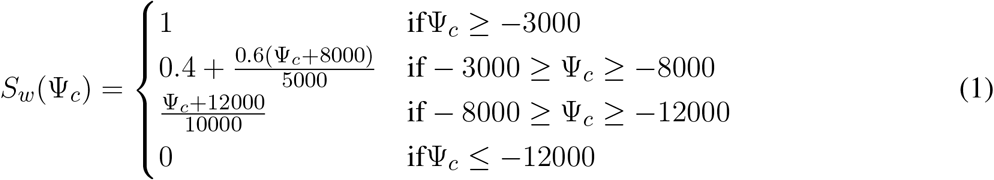

The precise relationship between leaf water potential and responses related to drought stress is difficult to establish but some estimates are available in the literature Acevedo et al. (1971); Tanguilig et al. (1987); Watts (1974). The water stress factor is translated to plant responses through transfer functions defined in the input file.

### 2.2 Photosynthesis

In order to have an accurate plant response to drought, the photosynthesis model we use should depend on solar irradiation as well as stomatal conductance. The Farquhar-Von Caemmerer-Berry model Farquhar and Caemmerer (1982); Farquhar et al. (1980); Von Caemmerer and Farquhar (1981) satisfies these conditions, is the most widely cited photosynthesis model and many published steady-state photosynthesis models are based on it or derived from it Morales et al. (2018). This model combines biochemical and irradiation dependent constraints and has been applied in a wide range of contexts Von Caemmerer (2013). For C3 plants the photosynthetic assimilation rate *A* in 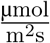 is equal to

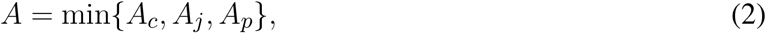

where *A*_*c*_ is the RuBisCo-limited (or carbon-limited) rate in 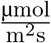, *A*_*j*_ the electron transport-limited (or light-limited) rate and *A*_*p*_ the phosphate-limited rate of carbon assimilation. We will assume that phosphate is not limiting so ignore *A*_*p*_ from here onward. *A*_*c*_ and *A*_*j*_ are defined as

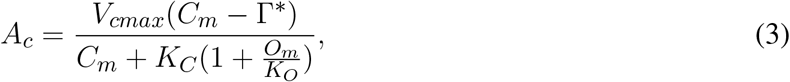

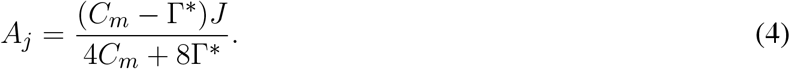

Here *V*_*cmax*_ is the maximum RuBisCo carboxylation rate in 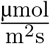, *K*_*C*_ and *K*_*O*_ are the RuBisCo Michaelis constants for CO_2_ and O_2_ in 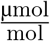 and Γ^∗^ is the CO_2_ compensation point without dark respiration (the leaf respiration that happens independently of photosynthesis) in ^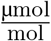^. *C*_*m*_ and *O*_*m*_ are the mesophyll CO_2_ and O_2_ concentrations in 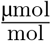 (see section 2.3). *J* is the potential electron transport rate in ^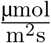^, which is defined as:

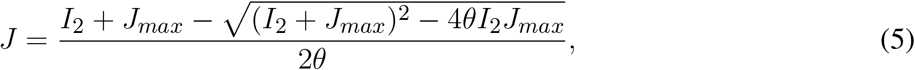

where *I*_2_ is the amount of energy in the light absorbed by photosystem II in 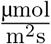, *J*_*max*_ is the maximum electron transport rate in 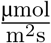 and *θ* is an empirical curvature factor. *I*_2_ is given by

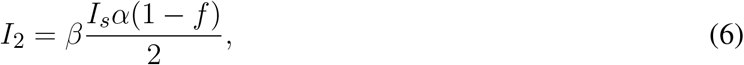

where *β* is a conversion factor in 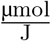, *I*_*s*_ is the solar irradiation in 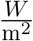, *α* the leaf absorptance and *f* a factor to correct for the spectral quality of the light.

The above equations are for C3 photosynthesis, we use the following expressions for C4 light-limited and carbon-limited photosynthesis rates Von Caemmerer (2000):

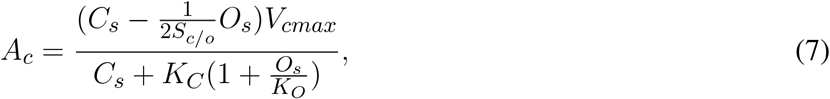

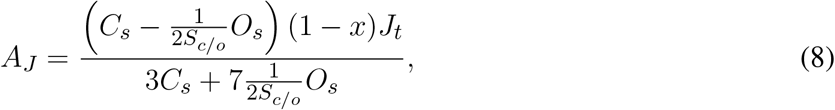

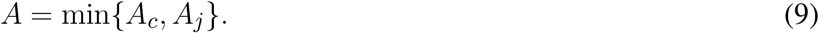

Here *C*_*s*_ and *O*_*s*_ are the bundle sheath carbon dioxide and oxygen concentrations in 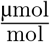, respectively and *S*_*c/o*_ is the RuBisCo specificity for CO_2_ relative to O_2_.

### 2.3 Leaf Gas Concentrations

Photosynthesis rates depend on the CO_2_ and O_2_ concentrations in mesophyll (C3 photosynthesis) or bundle sheath (C4 photosynthesis) cells. We formulate simple models for these concentrations. In the case of C3 photosynthesis, we model the mesophyll CO_2_ and O_2_ concentrations, *C*_*m*_ and *O*_*m*_ in 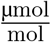, using the ordinary differential equations, which contain one term for the diffusion of CO_2_/O_2_ into/out of leaves through stomata, and one term for the depletion of CO_2_ and production of O_2_ because of photosynthesis and one term for the production of CO_2_ and depletion of O_2_ due to respiration. Assuming that diffusion of these gases is governed by Fick’s law Fick (1855), for the diffusion term we get the fluxes

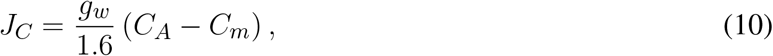

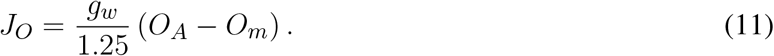

Here *g*_*w*_ is the stomatal conductance to water in 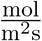, *C*_*A*_ is the atmospheric CO_2_ concentration in 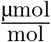 and *O*_*A*_ is the atmospheric O_2_ concentration in 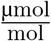. The ratios of the diffusivities in air of water, carbon dioxide and oxygen mean that 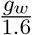 is the stomatal conductance to carbon dioxide and 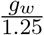 is the stomatal conductance to oxygen Haynes et al. (2016). The fluxes *J*_*C*_ and *J*_*O*_ both have the unit 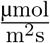. Dividing them by the thickness of the leaves *d*_*L*_ in metre we get the rate of change of gas molecules per unit volume, which has units 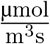. Since we denote concentrations in 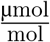, we want the rate of change per mol, instead of per unit volume. According to the ideal gas law, 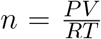, where *V* is a volume of gas in m^3^, *P* the pressure of that gas in Pa, *T* the temperature of that gas in K and *R* the ideal gas constant in 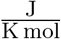. So by multiplying the fluxes by 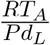 we get the rate of gas concentration change in our preferred units. Including terms for the photosynthetic assimilation rate *A* and the dark respiration rate *R*_*d*_, both in 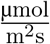, we get

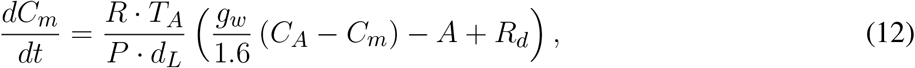

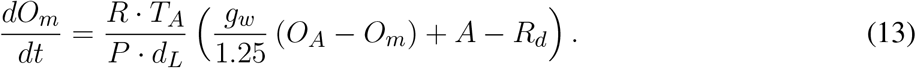

Here *A* is given by equation 2. The gas concentrations in leaves typically reach equilibrium conditions in less than a minute. Since typically *OpenSimRoot* timesteps are about a tenth of a day, we will make a quasi steady-state assumption. At steady-state,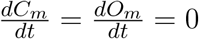 and equations 12 and 13 give

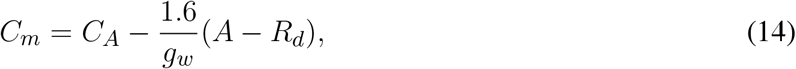

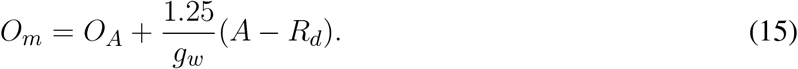

In C4 photosynthesis, the mesophyll exchanges gases with the atmosphere and CO_2_ is then transported to the bundle sheath. Therefore we model mesophyll and bundle sheath gas concentrations as connected reservoirs, using the differential equations:

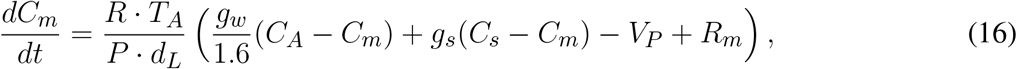

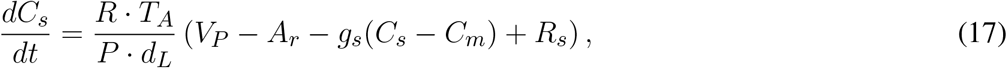

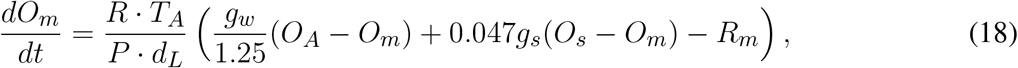

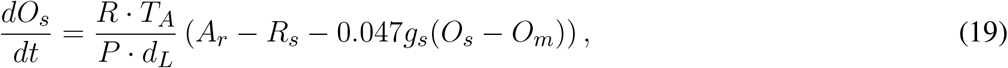

where *C*_*s*_ and *O*_*s*_ are the bundle sheath CO_2_ and *O*_2_ concentrations in 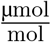, *R*_*m*_ and *R*_*s*_ are the respiration rates in mesophyll and bundle sheath cells in 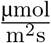, respectively, *g*_*s*_ is the conductance of the mesophyll-bundle sheath interface to CO_2_ in 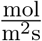, 0.047*g*_*s*_ is the mesophyll-bundle sheath interface conductance to O_2_ Von Caemmerer (2000), *V*_*P*_ is the PEP carboxylation rate in 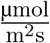 and *A*_*r*_ is the assimilation rate, not including dark respiration, in 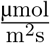. The equations concerning mesophyll concentrations have terms for gas exchange with both the atmosphere and the bundle sheath cells, a respiration term and a term for the PEP carboxylation, while the equations concerning bundle sheath concentrations have a term for the gas exchange with the mesophyll cells, a term for respiration and a term for photosynthesis. PEP carboxylation rates are given by

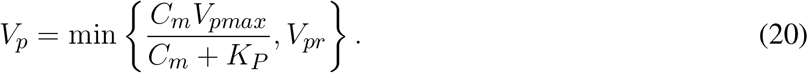

Here *V*_*pmax*_ in 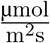 is the maximum PEP carboxylation rate, *K*_*P*_ in 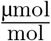 the Michaelis constant for PEP carboxylation and *V*_*pr*_ in 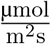 the PEP regeneration rate. As for C3 photosynthesis, gas concentrations reach equilibrium values very quickly as compared to the timescale relevant to *OpenSimRoot* and relative to changes in environmental conditions. Thus, we make a quasi steady-state assumption and we get:

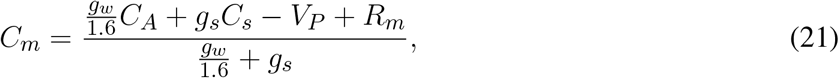

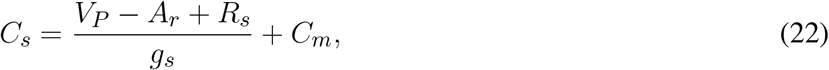

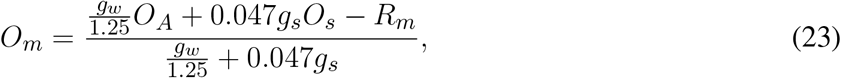

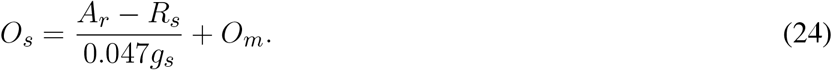

### 2.4 Stomatal Conductance

For the stomatal conductance to water *g*_*w*_, we use a slightly modified version of the Ball-Berry-Leuning model, described in Leuning (1995), which states that

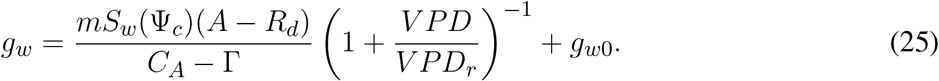

Here *m* is an empirical constant, *S*_*w*_(Ψ_*c*_) is the water stress factor defined in section 2.1, Γ in 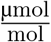 is the CO_2_ compensation point with dark respiration, *V PD* in Pa the vapour pressure deficit, *V PD*_*r*_ in Pa a reference vapour pressure deficit and *g*_*w*0_ in 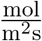the residual conductance. For C3 photosynthesis, Γ is given by

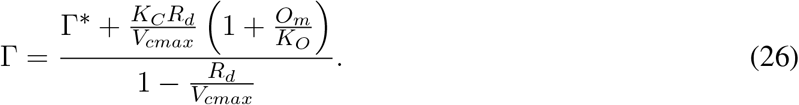

From Von Caemmerer (2000) we get the C4 expression for Γ, which is

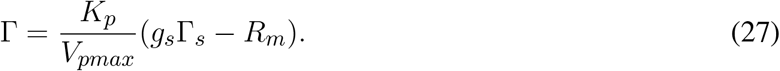

Here Γ_*s*_ in 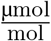 is the bundle sheath CO_2_ concentration at the compensation point, which is given by

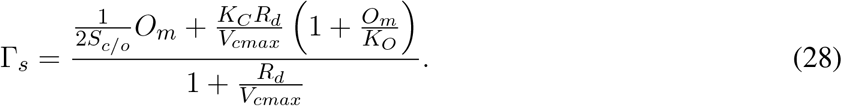

### 2.5 Leaf Temperature

Like many other aspects of plant development and function, photosynthetic assimilation rates are temperature dependent Duncan and Hesketh (1968); Lin et al. (2012). In order to make accurate predictions in the context of climate change, it is important that we can model the effects of elevated temperature and increasing atmospheric CO_2_ concentrations. This requires us to model the temperature dependence of the kinetic parameters that our models rely on Bowes (1991); Long (1991); McMurtrie and Wang (1993); Walcroft et al. (1997). For most parameters, we follow Von Caemmerer (2000) and use an Arrhenius function of the form

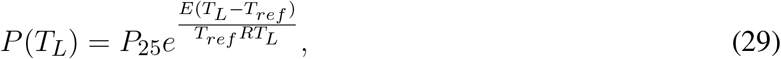

where *T*_*L*_ is the leaf temperature in K, *T*_*ref*_ the reference temperature, 25 ^°^C or 298.15 K, *P* (*T*_*L*_) is the value of the parameter at *T*_*L*_, *P*_25_ is the parameter value at 25 ^°^C (298.15K), *E* in 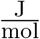 is the activation energy and *R* is the universal gas constant. We model the temperature dependence of Γ^∗^, *K*_*C*_, *K*_*O*_, *R*_*d*_, *V*_*cmax*_, *K*_*P*_ and 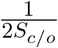 with equation 29. For other parameters, the peaked Arrhenius function of the form

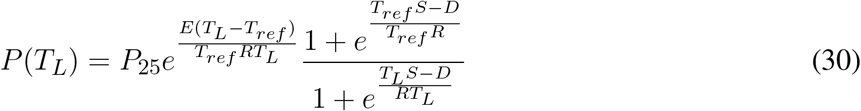

is used because it better represents the temperature dependence of that parameter. Here *D* in 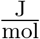 is the deactivation energy and *S* in 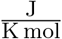 is called the entropy factor. We use equation 30 to model the temperature dependence of *J*_*max*_, *V*_*pmax*_ and *g*_*s*_. To calculate leaf temperature *T*_*L*_, we use an adaptation of the leaf energy balance as described in Nobel (2009). The terms that are described there as contributing to the leaf energy balance are: absorption of solar irradiation *A*_*s*_; absorption of infrared radiation from the surroundings *A*_*IR*_; emission of infrared radiation *e*_*IR*_; heat conduction and convection *H*_*c*_; heat loss due to evapotranspiration *H*_*t*_; photosynthesis; metabolic processes.

The contribution of photosynthesis and metabolic processes to the leaf energy balance is typically on the order of a few Watts, while other terms such as the solar irradiation are several hundreds of Watts, so to simplify the model we omit the photosynthesis and metabolic processes from the energy balance. Due to the low specific mass of leaves (approximately 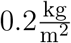), their specific heat is low and a net energy balance of a few Watts is enough to increase the temperature by a degree Celsius in less than 5 minutes. Because of this, we assume the leaf is in steady state and the energy balance *E* is zero:

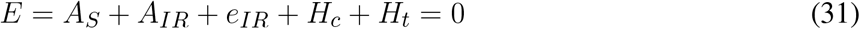

The absorption of solar irradiation per unit area *A*_*S*_ in 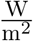 is equal to

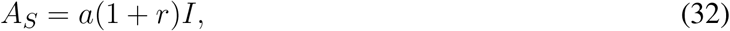

where *a* is the absorptance of the leaf over the whole solar spectrum, *r* is the fraction of solar irradiation reflected by the surroundings and *I* in 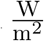 is the solar irradiation. Assuming there are no nearby objects that reflect a lot of sunlight towards the field in which we are modelling crops and using a symmetry argument (for each photon reflected from another leaf, we can assume the leaf reflects a photon itself), we set *r* = 0. So that

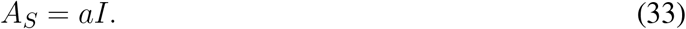

Using the Stefan-Boltzmann law, the absorption of infrared radiation per unit area *A*_*IR*_ in 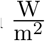 is

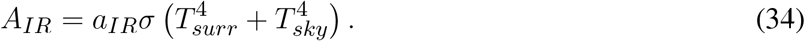

Here *a*_*IR*_ is the infrared radiation absorptance of the leaves, *σ* in 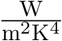 is the Stefan-Boltzmann constant, *T*_*surr*_ in K is the temperature of the surroundings and *T*_*sky*_ in K is the effective temperature of the sky. Here we assume that the underside of the leaves absorb infrared radiation coming from the surroundings while the upper side absorbs infrared radiation coming from the sky. Note that *T*_*sky*_ is not the actual temperature of the sky, but the temperature a blackbody that emitted as much radiation as the sky would have. We assume that

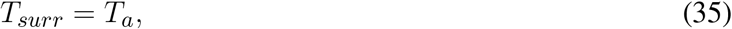

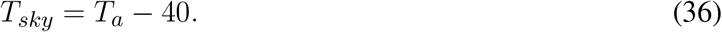

A blackbody with absorptance *a* will also have emissivity *a* so the emission of infrared radiation by the leaves, *e*_*IR*_ in 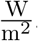, is equal to

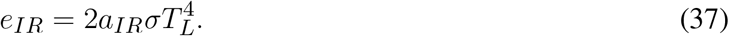

The factor 2 in this equation 37 comes from the fact that the leaves emit infrared radiation from both sides. The heat loss due to conduction into the air boundary layer around leaves and then convection away from the leaves, *H*_*c*_ in 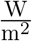, is equal to

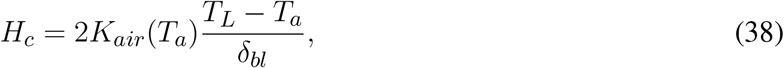

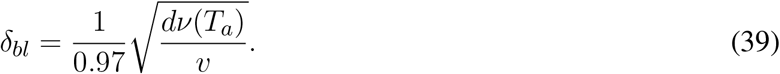

Here *K*_*air*_(*T*_*a*_) in 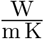 is the thermal conductivity of the air, *δ*_*bl*_ in m the thickness of the boundary layer, *d* in m the characteristic length of the leaf, *ν*(*T*) in 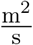 the kinematic viscosity of the air and *v* in 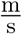 the wind speed. The factor 2 in equation 38 is because heat is conducted away from both sides of the leaf. For the range of air temperatures we are concerned with, 0 to 50 ^°^C (273.15 to 323.15 K), we use, from Nobel (2009), the following approximations

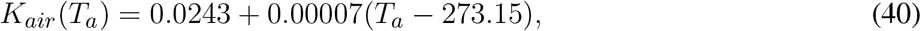

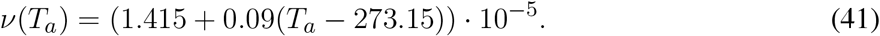

The heat loss due to evapotranspiration, *H*_*t*_ in 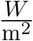, is equal to

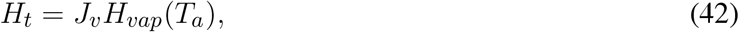

where *J*_*v*_ in 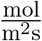 is the transpiration rate and *H*_*vap*_(*T*_*a*_) in 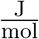 is the heat of vaporisation of water, which is equal to

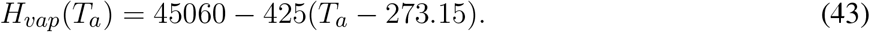

Putting this together and setting the leaf energy balance equal to zero we obtain

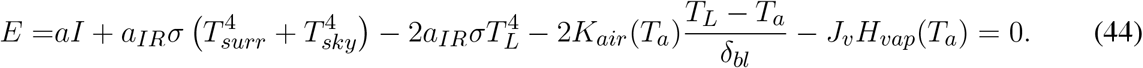

This is a quartic equation in *T*_*L*_, the leaf temperature, which we solve numerically using the Newton-Raphson method. As our initial guess we take *T*_*a*_, the atmospheric temperature, which should be somewhat close to *T*_*L*_.

### 2.6 Scaling from Leaf to Canopy

The photosynthesis, leaf gas concentration, stomatal conductance and leaf temperature models described in previous sections are all models for individual leaves. However, in a canopy, not every leaf is subjected to the same conditions. Some leaves are in direct sunlight, while others will be partially or completely shaded by other leaves. A number of different models have been proposed to scale up from the leaf scale to the canopy scale. The simplest, big leaf models, approximates the entire canopy as a single leaf Amthor (1994); Lloyd et al. (1995). These models tend to overestimate photosynthesis rates and require empirical extinction factors to make them produce accurate results. More sophisticated models use multi-layer approaches, which divide the canopy up in layers, applying leaf-scale models to each separately Chang et al. (2018); de Wit (1965); Duncan et al. (1967); Lemon et al. (1971); Waggoner and Reifsnyder (1968). We use a third approach, the sun/shade model described in De Pury and Farquhar (1997), which divides the canopy up into sunlit and shaded leaf area fractions, producing results with similar accuracy to the multi-layer model De Pury and Farquhar (1997) while also being relatively simple. This is the model we chose to represent our canopy. In this model the sunlit and shaded leaf area fractions are calculated each time step based on leaf area index and the angle of the sun with respect to the canopy. We integrate this with the previously described models by calculating stomatal conductance, leaf gas exchange, leaf temperature and photosynthesis rates separately for the shaded and sunlit leaf areas.

### 2.7 Nitrogen limitations on photosynthesis

Leaf nitrogen content is an important factor determining photosynthetic assimilation rates Sinclair and Horie (1989). Using existing *OpenSimRoot* models for minimum and optimum nutrient concentrations, which have to be specified for every plant organ, and the total plant nitrogen content, we estimate leaf nitrogen content leaf_*N*_ :

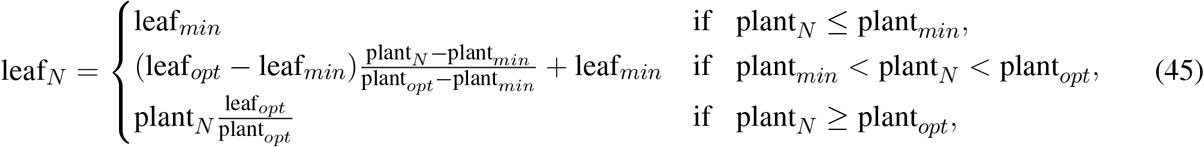

where leaf_*min*_ and leaf_*opt*_ are the minimal and optimal leaf nitrogen contents, plant_*min*_ and plant_*opt*_ are the minimal and optimal plant nitrogen contents and plant_*N*_ is the total plant nitrogen content.

We assume the electron transport rate *J*_*max*_ and carboxylation rate *V*_*cmax*_ are limited by leaf nitrogen content through the folowing equations:

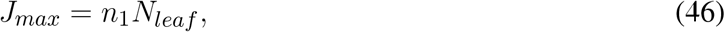

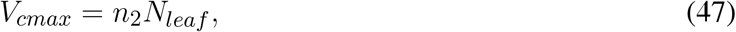

where *n*_1_ and *n*_2_ are proportionality constants Kull and Kruijt (1998) and *N*_*leaf*_ is the leaf nitrogen concentration (not content as in equation 45). By substituting *J*_*max*_ into equation 5 and *V*_*cmax*_ in equation 3 (or equation 7 in the case of C4 photosynthesis), nitrogen limitations are taken into account.

### 2.8 Carbon Reserves

*OpenSimRoot* contains a relatively simple model for the accumulation and degradation of starch carbon reserves. From the available carbon, *C*_*A*_ in 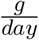, from photosynthesis and the seed, carbon costs such as *C*_*C*_ *in* 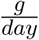, are first deducted. Potential growth rates of different plant organs together determine the carbon sink for growth, *C*_*G*_ in 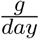. Any remaining carbon after allocating to growth is added to the carbon reserves (a pool representing the carbon in starch and other metabolites). The allocation rate of carbon to reserves is denoted by *C*_*R*_ in 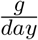 (note that if carbon is remobilized from the reserves, *C*_*R*_ *<* 0), while the current amount of carbon in the reserves is denoted by *C*_*P*_ in *g*. So in most cases

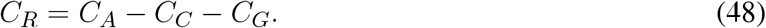

If not enough carbon is available from photosynthesis and the seed for carbon costs and growth, so if *C*_*A*_ < *C*_*C*_ + *C*_*G*_, carbon from the reserves is remobilised. The plant cannot remobilize more than half of the current reserves per day, so 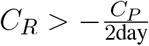. If carbon from photosynthesis, the seed and reserves together is not enough to match the carbon costs and growth, so if *C*_*A*_ − *C*_*R*_ < *C*_*C*_ + *C*_*G*_, the allocation of carbon to growth will be less than the amount required for potential growth. The actual amount of carbon available for growth, *C*_*H*_, will then be *C*_*H*_ = *C*_*A*_ *C*_*R*_ *C*_*C*_ < *C*_*G*_. This will lead to a reduction in growth rates, thus controlling growth through carbon availability.

This carbon reserve allocation model works well with a photosynthesis model based on light-use efficiency which averages out photosynthesis over the full 24 hours in a day. However, since we are now adding models for simulating the day-night cycle, we need to allow for carbon to be stored during the day and then released during the night. The model has to satisfy these important requirements:

- Carbon remobilisation from the reserves at night should be high enough to allow for uninterrupted growth if water and nutrients are plentiful.
- Allocation of carbon to the reserves during the day should be high enough such that there is enough in reserves to match the non-growth carbon costs during the night.
- If photosynthesis rates over a 24-hour period are lower than the carbon needed for maintenance and growth then growth rates should be reduced.

Analysis of Arabidopsis carbon reserves under photoperiods of different length showed that starch reserves were remobilised at fairly constant rates which depended on the length of the photoperiod Sulpice et al. (2014). It was also found that starch accumulation was more rapid in shorter photoperiods. Based on this paper and our requirements, we implement the following modifications to the model of carbon reserves.

- At night, the maximum rate of starch remobilisation is determined at the start of the night and set such that at most 95% of the reserves are remobilised during the night. 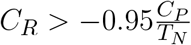 where *T*_*N*_ is the duration of the night.
- During the day, if carbon reserves at the end of the day are expected to be below 110% of the total carbon needed during the night, growth is reduced. More precisely, if *C*_*P*_ + *C*_*R*_*T*_*S*_ ≤ 1.1*C*_*C*_*N*, where *T*_*S*_ is the time until sunset, then *C*_*R*_ is bounded from below by min 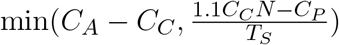.

### 2.9 *OpenSimRoot* Root Finder

At any given time, in order to calculate the photosynthetic assimilation rates, we have to find a solution to the system of of equations 2, 3, 4, 14, 15 and 25 (in the case of C4 photosynthesis this is 21, 22, 23, 24, 20, 7, 8, 9 and 25). It has been shown that when solving the coupled photosynthesis and stomatal conductance equations using a fixed point method there is no convergence in many cases Sun et al. (2012). In order to guarantee convergence and arrive at the correct solution in that case, it has been suggested to use the Newton-Raphson method Sun et al. (2012). Ideally, we would use a multivariate Newton-Raphson method to solve the above system of equations. However, this is only possible within the *OpenSimRoot* engine if the whole system of equations and relevant solutions are solved within a single submodel. Doing this would make it much more complicated for future users to modify the equations, choose different models for one of the relevant state variables or add new models coupled to the equations above, which is what the modular structure of the *OpenSimRoot* engine was designed to facilitate. By keeping models separate, we allow users to pick and choose between different models for variables like photosynthesis, leaf gas exchange, leaf temperature or stomatal conductance. Because of this, we have implemented a root finder based on the Newton-Raphson method which works within the constraints of the *OpenSimRoot* engine. Suppose we have a system of equations

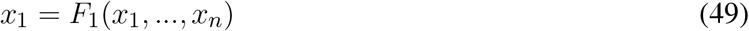

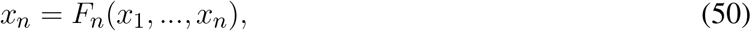

where *x*_1_ could for example be the stomatal conductance and *F*_1_ equation 25, the expression for stomatal conductance. We start from initial values *x*_1,0_, …, *x*_*n*,0_. The root finder will then use the Newton-Raphson method to find an *x*_*n*,∗_ for which

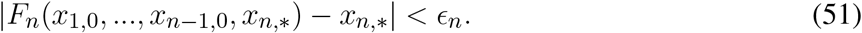

Here *ϵ*_*n*_ is a tolerance. Then, we want to find a *x*_*n*−1,∗_ which satisfies

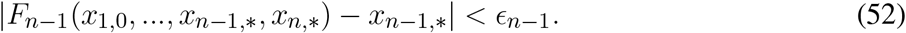

We can again do this using the Newton-Raphson method but at each step will update *x*_*n*,∗_. More concretely, the root finder does one step of the Newton-Raphson method to find *x*_*n*−,1_ and finds a new *x*_*n*,∗_ that now satisfies

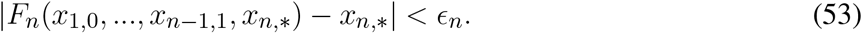

After a number of steps we find an *x*_*n*−1,∗_ which satisfies

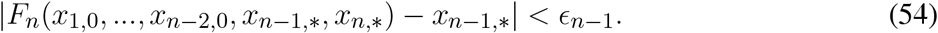

Now we use the same process to find *x*_*n* − 2,∗_ which satisfies

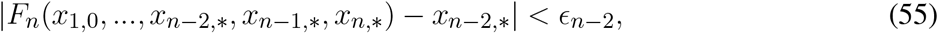

again updating *x*_*n* − 1,∗_ and *x*_*n*,∗_ at each iteration step. Following this iterative procedure for all variables we eventually find *x*_1,∗_, *x*_2,∗_, …, *x*_*n*,∗_ which satisfy

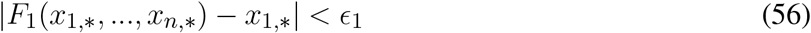

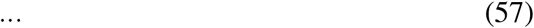

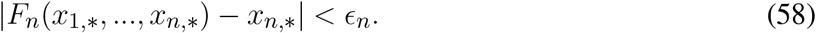

This is the simultaneous solution to the system of equations making up the photosynthesis model. If there is no convergence in 40 steps, the value from the previous timestep will be used (in case this happens on the first timestep, a predetermined default value will be used).

### 2.10 Simulation setup

In this paper, we test the utility of several variants of the “Steep, Cheap, and Deep” (SCD) ideotype against a reference root phenotype of maize under rainfed and drought conditions in three distinct maize-producing pedo-climatic environments. The SCD phenotypes are different from the reference phenotype in 4 phene states:

- The number of axial roots from different classes (i.e., seminal, nodal, and brace) is reduced from 89 to 45.
- The angles of all axial root classes (i.e., seminal, nodal, and brace) are 20 degrees steeper compared to the reference phenotype.
- The lateral root branching density (LRBD) is reduced by 50%.
- The maximum root cortical aerenchyma (RCA) volume is increased from 20% to 40%.

In addition to simulating the reference and SCD phenotype, we simulated every combination of the reference and SCD values for the above 4 phenes resulting in 16 different combinations. We will refer to these different phenotypes as follows:

- All phenes have the reference value: Reference phenotype.
- All phenes have the reference value except one which has the SCD value: The phene change from the reference value (so “Fewer axial roots”, “steeper axial roots”, “reduced lateral root branching density”, “higher root cortical aerenchyma”).
- Two phenes with the reference value, two with the SCD value: The phene values which are not the reference value (so for example: “Fewer, steeper axial roots”).
- One phene with the reference value, three with the SCD value: SCD + the phene value which has the reference value (so for example: “SCD + more axial roots” if the first phene from the list above has the reference value).
- All four phenes have the SCD value: Steep, cheap, deep (SCD).

We test the utility of these 16 root phenotypes under rainfed and terminal drought conditions in three different pedo-climatic environments. A single maize plant is simulated representing an individual within a monoculture stand with a between-row spacing of 50 cm, within-row spacing of 25 cm, and soil depth profile of 150 cm, corresponding to the planting density of 8 plants m-2 and sowing depth of 5 cm. Roots from neighboring plants are simulated by mirroring the roots at the boundary back into the column. To accurately mimic the rainfed maize agrosystem, actual soil and climate datasets for three different maize growing locations, namely Boone County (Iowa, USA - 42^°^49.4’N 93^°^43’45.9”W), Zaria (Kaduna, Nigeria - 11^°^1’N 7^°^37’E), and Tepatitlán (Jalisco, Mexico - 20^°^52’N 102^°^43’W) are used. For simplicity, these sites are referred to as Iowa, Zaria, and Jalisco. The soils in these sites have been identified as *Mollisol – Udoll* (Iowa), *Alfisol – Ustalf* (Zaria), and *Andosol – Vitrand* (Jalisco). Soil profile data, including soil texture, soil bulk density, and relative water content at -33 and -1500 kPa, from the ISRIC soil hub database (https://data.isric.org/) is used for Iowa and Jalisco, and for Zaria a published dataset is used Beah et al. (2020). These soil profile datasets are implemented to estimate the corresponding van Genuchten parameters using the online freeware Rosetta3 (https://www.handbook60.org/rosetta/). The weather data for the year 2019 over the main growing season for all three locations come from the *POWER LaRC* database (https://power.larc.nasa.gov/data-access-viewer/). For the drought scenarios, we simulate a terminal vegetative stage drought by setting precipitation rates to zero after a certain number of weeks. This time point is chosen such that the reference phenotype would see a reduction in shoot dry weight of approximately 50% as compared to the rainfed scenario. For Iowa, the terminal vegetative drought starts on day 21, while for the Zaria and Jalisco it starts on day 28.

Instructions on how to acquire the correct executable and reproduce the simulations done in this study can be found at GitLab-OpenSimRoot v2.See Schäfer et al. (2022b) for a comprehensive introduction on running *OpenSimRoot*. Supplementary Information 1 contains an input file with all parameters corresponding to the new capabilities.

### 2.11 Estimating phene interactions

The actual (predicted) shoot biomass in response to the drought of each integrated SCD phenotype is compared to their corresponding expected shoot biomass response to quantify the interactions among the root phenes. The expected shoot biomass for each SCD phenotype is derived using the additive null model Côté et al. (2016) and the shoot biomass response of the reference maize root phenotype. The shoot biomass responses of the individual phene states are calculated by subtracting the reference shoot biomass from each SCD phenotype that differs in just one phene state. The resulting shoot biomass responses of individual phene states are then added to the reference shoot biomass to get the expected additive shoot biomass corresponding to a combination of phene states. The variance of the expected additive shoot biomass is calculated by summing the variances of the relevant quantities (the variance of a sum of distributions is the sum of variance). From this variance we calculate the standard error by taking 5 as the population size, matching the number of repetition simulations we run for each set of inputs. The actual response greater than, equal to, and less than the expected response respectively corresponds to the synergistic, additive, and antagonistic relationship between the root phenes. More details on this can be found in Ajmera et al. (2022); Ma et al. (2001).

## 3 RESULTS

The results highlighted in this study demonstrate the new capabilities that mechanistically capture processes underlying shoot physiology in *OpenSimRoot_v2* (Figure 1). The impact of maize root phenotypes on shoot dry weight varied in response to both pedo-climatic conditions and drought status (Figure 2). In the rainfed Iowa environment, the reference phenotype and the high aerenchyma phenotype had the greatest shoot dry weight with just under 25 g. In response to drought in Iowa, the shoot dry weight of the reference phenotype was reduced to 13 g. However, phenotypes with fewer, shallow axial roots and reduced lateral root branching density had the greatest shoot dry weights under drought in Iowa (approximately 20 g). Steep axial roots were not associated with high shoot dry weight in Iowa drought conditions and the amount of RCA had little effect on shoot dry weight in these conditions.

**Figure 2.**
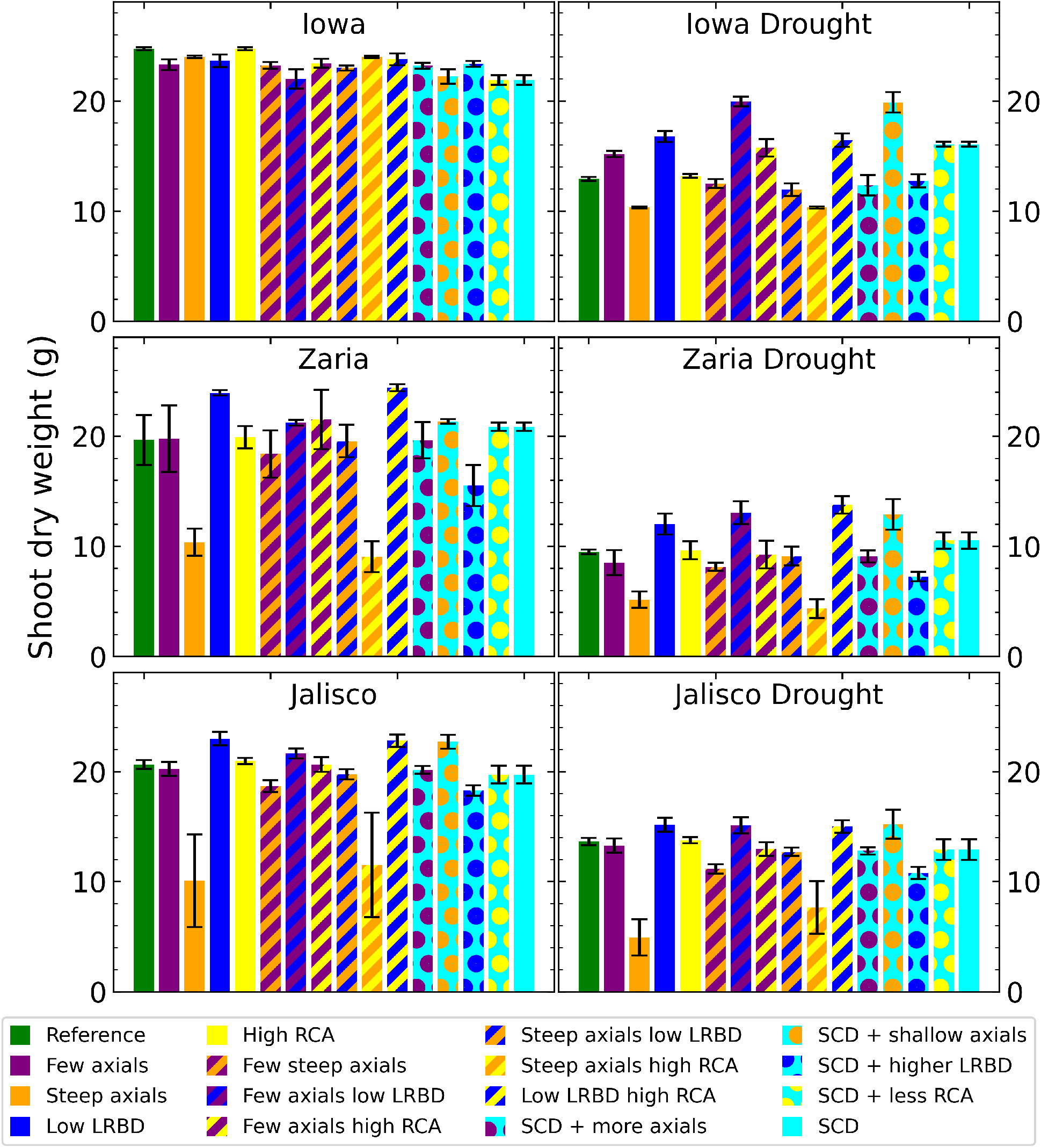
Shoot dry weight of all the maize phenotypes considered after 42 days in the 6 different environments. The error bars indicate standard deviation. LRBD means lateral root branching density, RCA means root cortical aerenchyma and SCD refers to the steep, cheap, deep phenotype.

In Zaria, the reference phenotype had intermediate performance in both the rainfed and drought conditions, with 19.75 and 9.5 g of shoot dry weight. In rainfed conditions, two phenotypes with shallow axial roots and abundant aerenchyma had shoot dry weights smaller than 10 g, indicating that rainfed Zaria conditions were more stressful than the rainfed Iowa conditions. In Zaria, reduced lateral branching density was associated with increased shoot dry weight in combination with most other phenes under drought conditions. In contrast to Iowa, reducing the number of axial roots without any other changes in root phenotypes was slightly detrimental in terms of shoot biomass gain in response to drought in Zaria. In Jalisco, both the reference and SCD phenotypes were near the greatest shoot dry weight in both rainfed and drought conditions. The phenotypes with reduced lateral branching density (LRBD), few axials plus reduced LRBD, reduced LRBD plus abundant aerenchyma, and shallow axial roots were the best performers in both environments. Except for rainfed Iowa, across all the environments the phenotypes with steep axial roots plus at either level of aerenchyma had the smallest shoot dry weight.

Under rainfed conditions, the SCD phenotype had significantly smaller root carbon costs (structural carbon, respiration, and exudation) than the reference phenotype, with a reduction of approximately 30% in Zaria (Figure 3). Under drought, the difference between root carbon costs of the reference and SCD phenotypes decreased in the Iowa and Zaria environments and stayed the same in the Jalisco environment. Decreasing the axial root number reduced root carbon costs the most in all environments except the Jalisco drought environment. The effect of other single phene changes depend more on the environment. Phenotypes with few axial roots combined with abundant aerenchyma and large lateral root branching density had small root carbon costs in most environments.

**Figure 3.**
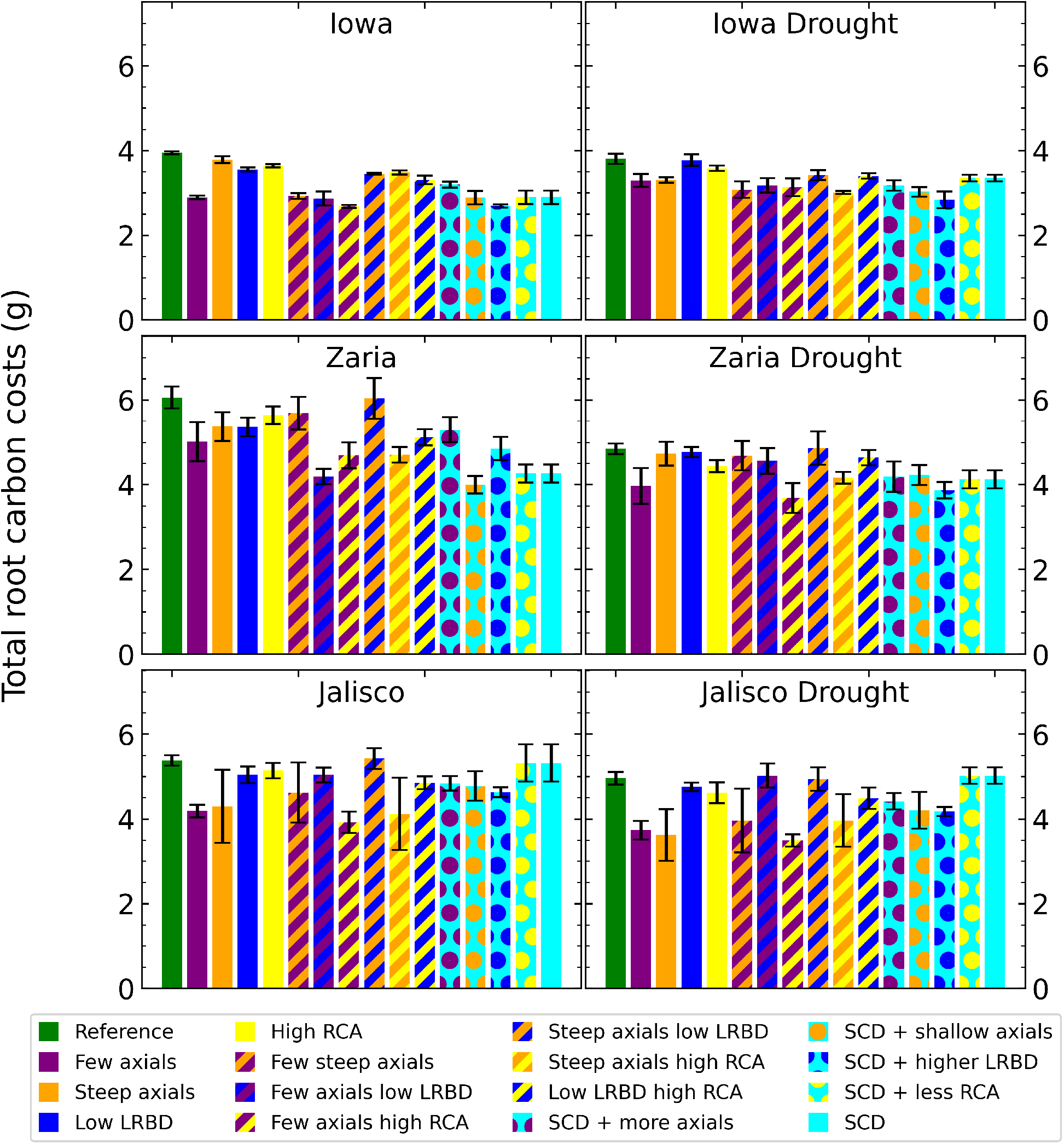
Total carbon invested in roots (biomass, exudation and respiration) of all the maize phenotypes considered after 42 days in the 6 different environments. The error bars indicate standard deviation. LRBD means lateral root branching density, RCA means root cortical aerenchyma and SCD refers to the steep, cheap, deep phenotype.

The carbon use efficiency (water acquired per unit of carbon invested in roots) of the SCD phenotype was greater than that of the reference phenotype in all environments, (although the difference was very small in Jalisco (Figure 4). From the individual changes, fewer axial roots, reduced lateral root branching density, and greater aerenchyma formation improve carbon use efficiency, while steeper axial roots reduce carbon use efficiency. In Iowa, the carbon use efficiency was greater than in the other two environments by about 0.5 liters per gram of carbon.

**Figure 4.**
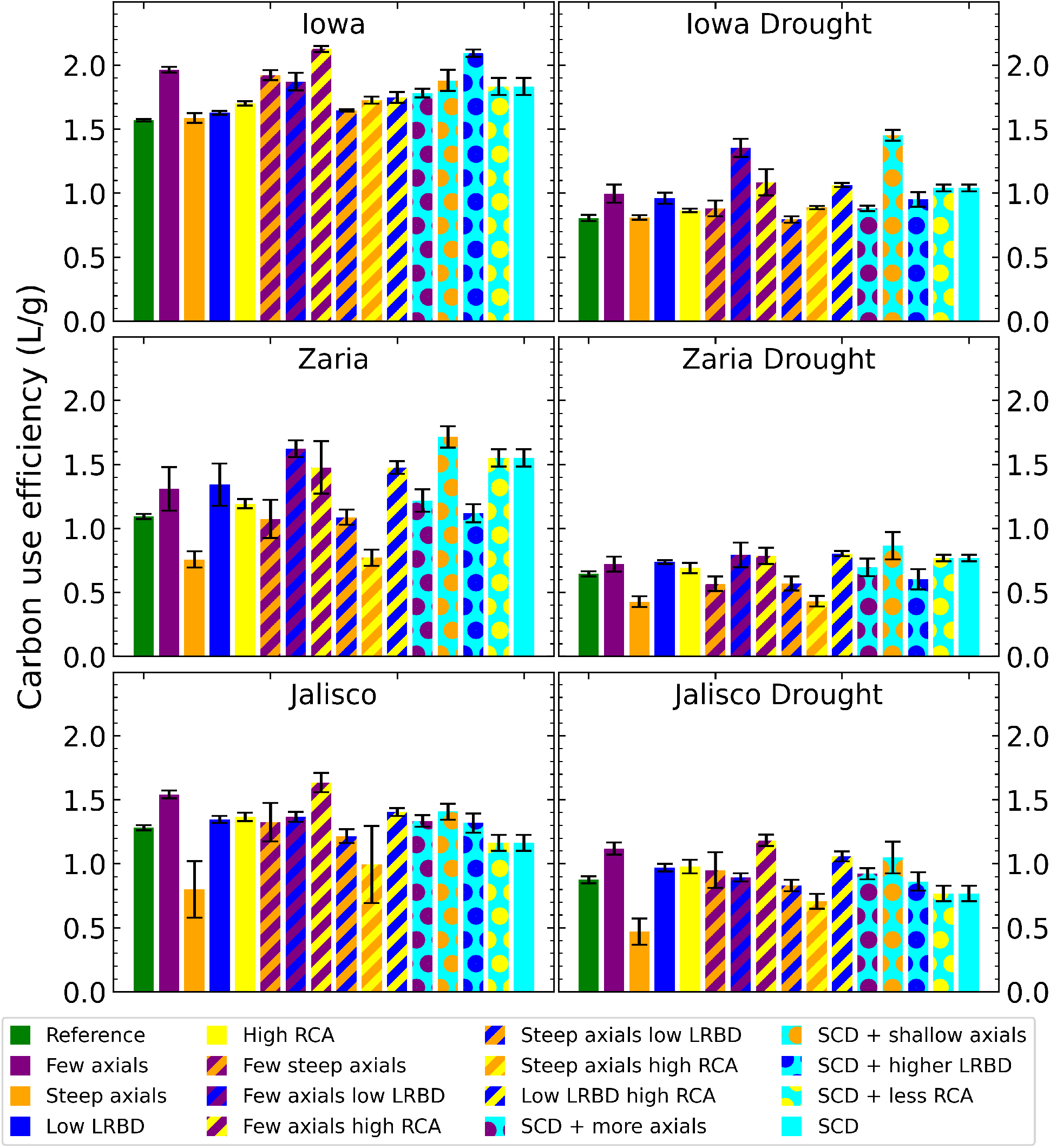
Carbon use efficiency (water uptake per unit of carbon invested in roots) of all the maize phenotypes considered after 42 days in the 6 different environments. The error bars indicate standard deviation. LRBD means lateral root branching density, RCA means root cortical aerenchyma and SCD refers to the steep, cheap, deep phenotype.

The SCD phenotype, with either aerenchyma level, has the largest fraction of root length deeper than 50 cm (Figure 5). In Iowa, 1% or less of roots in the reference phenotype were deeper than 50 cm, versus more than 8% for the SCD phenotype. In Zaria, 10% of the root length was below 50 cm for the reference phenotype, which increased to 20% in the rainfed environment and 24% in the drought environment for the SCD phenotype. In Jalisco, about 17% of root length was below 50 cm for the reference phenotype, which increased to ca. 30% for the SCD phenotype. Reducing axial root number lead to a bigger increase in deep roots than making the axial root angle steeper, as does reducing lateral root branching density in the majority of cases. Aerenchyma formation had little effect on the fraction of deep roots.

**Figure 5.**
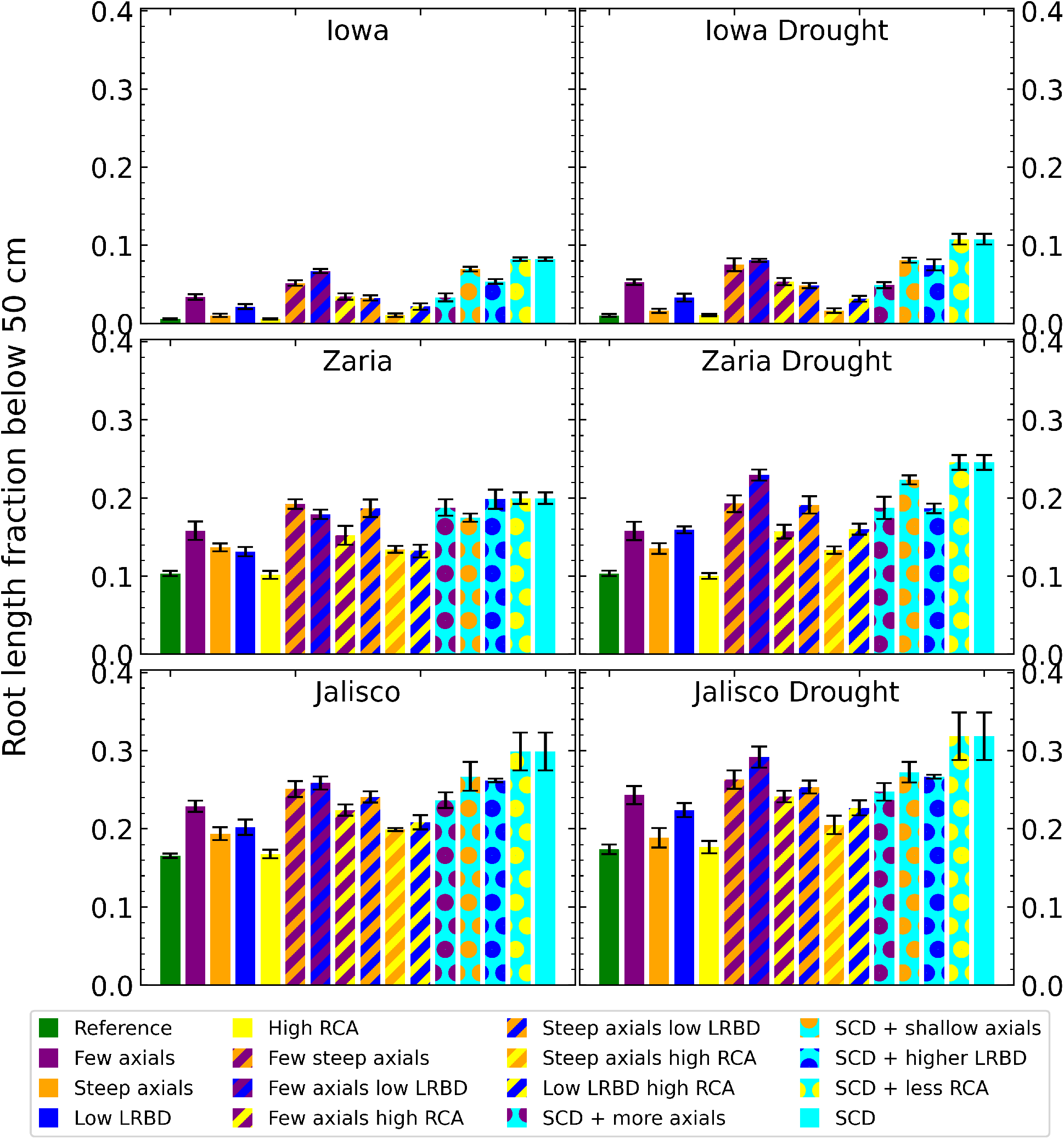
Fraction of the total root length below 50 cm deep in the soil of all the maize phenotypes considered after 42 days in the 6 different environments. The error bars indicate standard deviation. LRBD means lateral root branching density, RCA means root cortical aerenchyma and SCD refers to the steep, cheap, deep phenotype.

There were major differences in the depth profiles of soil penetration resistance encountered by roots among the 6 environments (Figure 6). In general, rainfed environments had less soil penetration resistance than drought environments: the difference between them was often several thousand kPa. In rainfed Iowa soil, penetration resistance sharply increased between 20 and 45 cm. Under drought, Iowa soil had the greatest penetration resistance of all the soils, peaking at more than 14000 kPa at 45 cm. The soil penetration resistance of the rainfed Zaria soil increased from around 900 kPa at the soil surface to 5000 kPa at 150 cm. In the Zaria drought environment, the soil penetration resistance was greater at every depth and reached its maximum values between 50 and 80 cm deep, after which it declined again. In the Jalisco rainfed environment, the reference phenotype encountered greater soil penetration resistance than the SCD phenotype, the soil penetration resistance encountered by the reference phenotype is similar to what both phenotypes encounter in the Jalisco drought environment.

**Figure 6.**
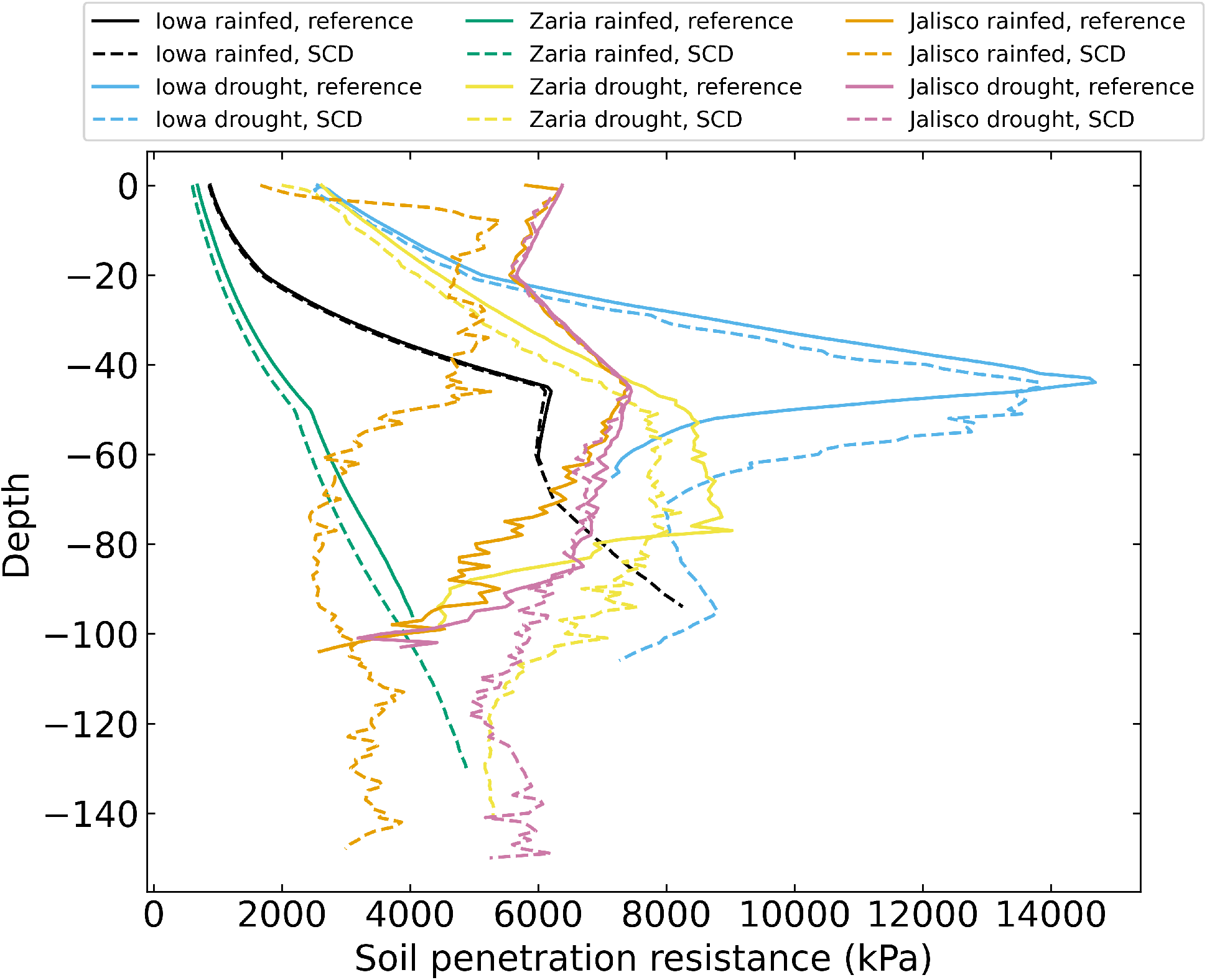
Soil penetration resistance felt by the roots on day 42 for 12 individual simulations, averaged over 1 cm layers.

Plants with the SCD phenotype had more water near their roots at most depths than plants with the reference phenotype (Figure 7). In the rainfed Iowa and Zaria environments, there was more water near the roots in the topsoil than there was in deeper soil layers. In drought environments, there was less water near the roots than in rainfed environments and there was relatively little water near the roots in the topsoil and more in deeper soil strata.

**Figure 7.**
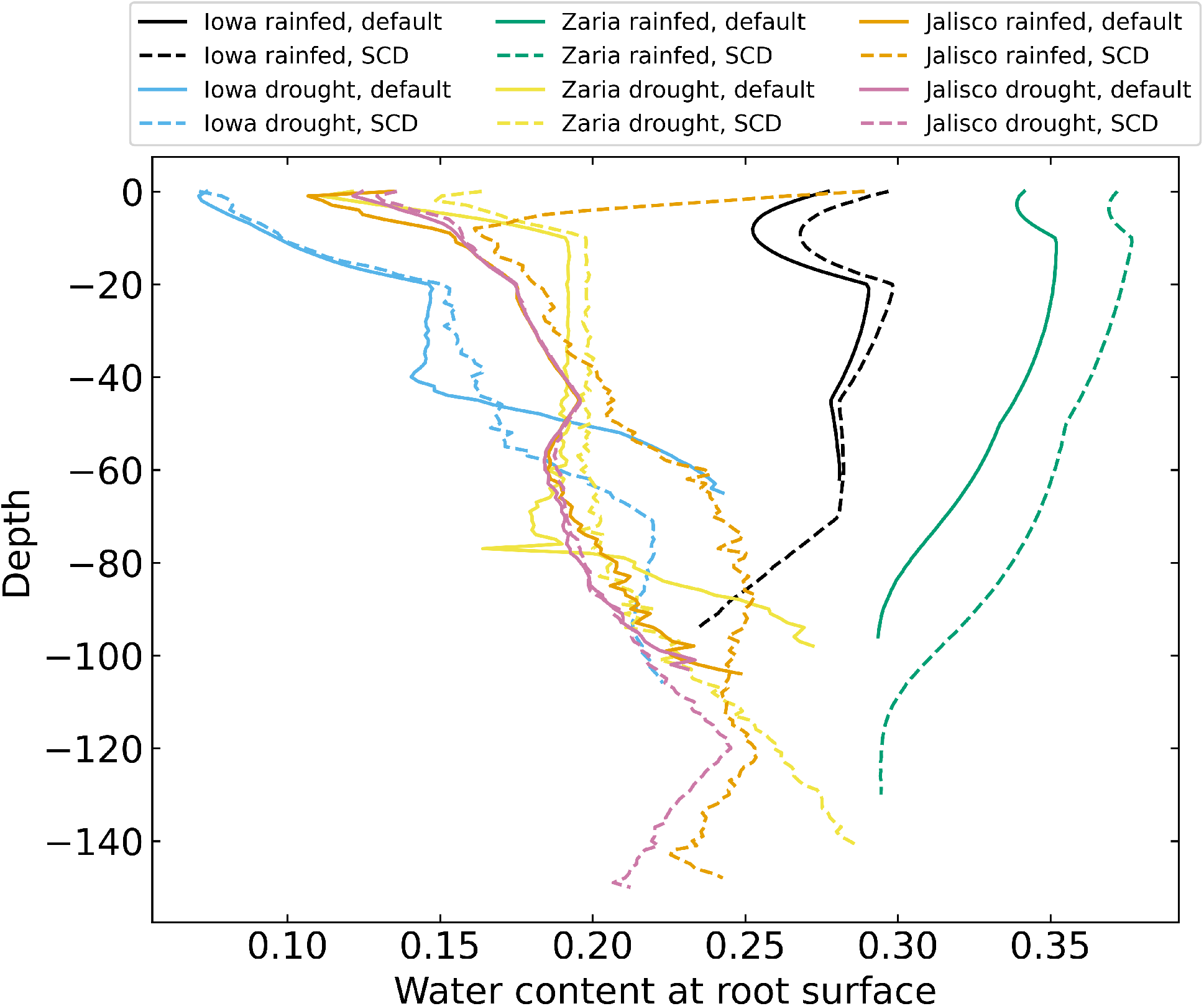
Volumetric water content near the roots on day 42 for 12 individual simulations, averaged over 1 cm layers.

The SCD phenotype had significantly greater carbon deposition (carbon in the form of root biomass or exudates) below 50 cm in the soil than the reference phenotype in all 6 environments (Figure 8). In Iowa, the SCD phenotype deposited approximately 8 times more carbon deep in the soil than the reference phenotype under both rainfed and drought conditions. In the Zaria soil, the SCD phenotype deposited 1.6 times more deep carbon in the rainfed case and 2.2 times more in the drought case, while in the Jalisco soil, the difference was a factor 2 in both cases. Besides the SCD phenotype, a few other phenotypes had similar or even greater carbon deposition. The performance of the SCD phenotypes with less aerenchyma formation or shallow axials were similar in most environments. This was similar to the performance of the phenotypes with few steep axial roots and those with low lateral root branching density in several environments.

**Figure 8.**
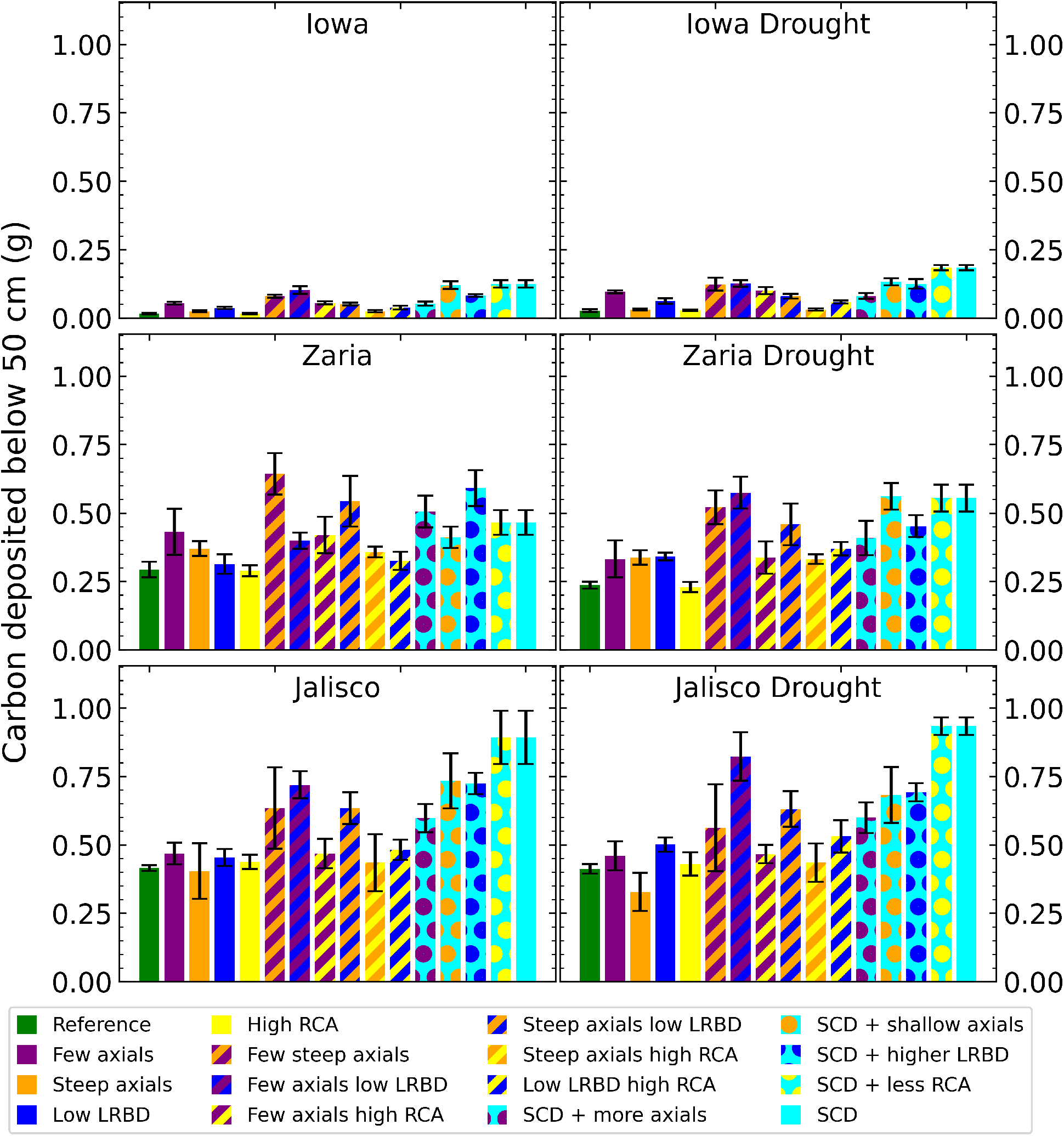
Carbon depositions below 50 cm deep in the soil of all the maize phenotypes considered after 42 days in the 6 different environments. The error bars indicate standard deviation. LRBD means lateral root branching density, RCA means root cortical aerenchyma and SCD refers to the steep, cheap, deep phenotype.

Except in the rainfed Iowa environment, there was at least one SCD variant phenotype that had both greater shoot dry weight and greater carbon deposition below 50 cm than the reference phenotype (Figure 9). In the rainfed Iowa environment, the reference phenotype had the greatest shoot dry weight and least deep soil carbon deposition while the SCD phenotype had one of the greatest deep soil carbon depositions and smallest shoot dry weights. In almost all environments, steeper axial roots by themselves significantly reduced shoot dry weight for a relatively small increase in deep soil carbon deposition.

**Figure 9.**
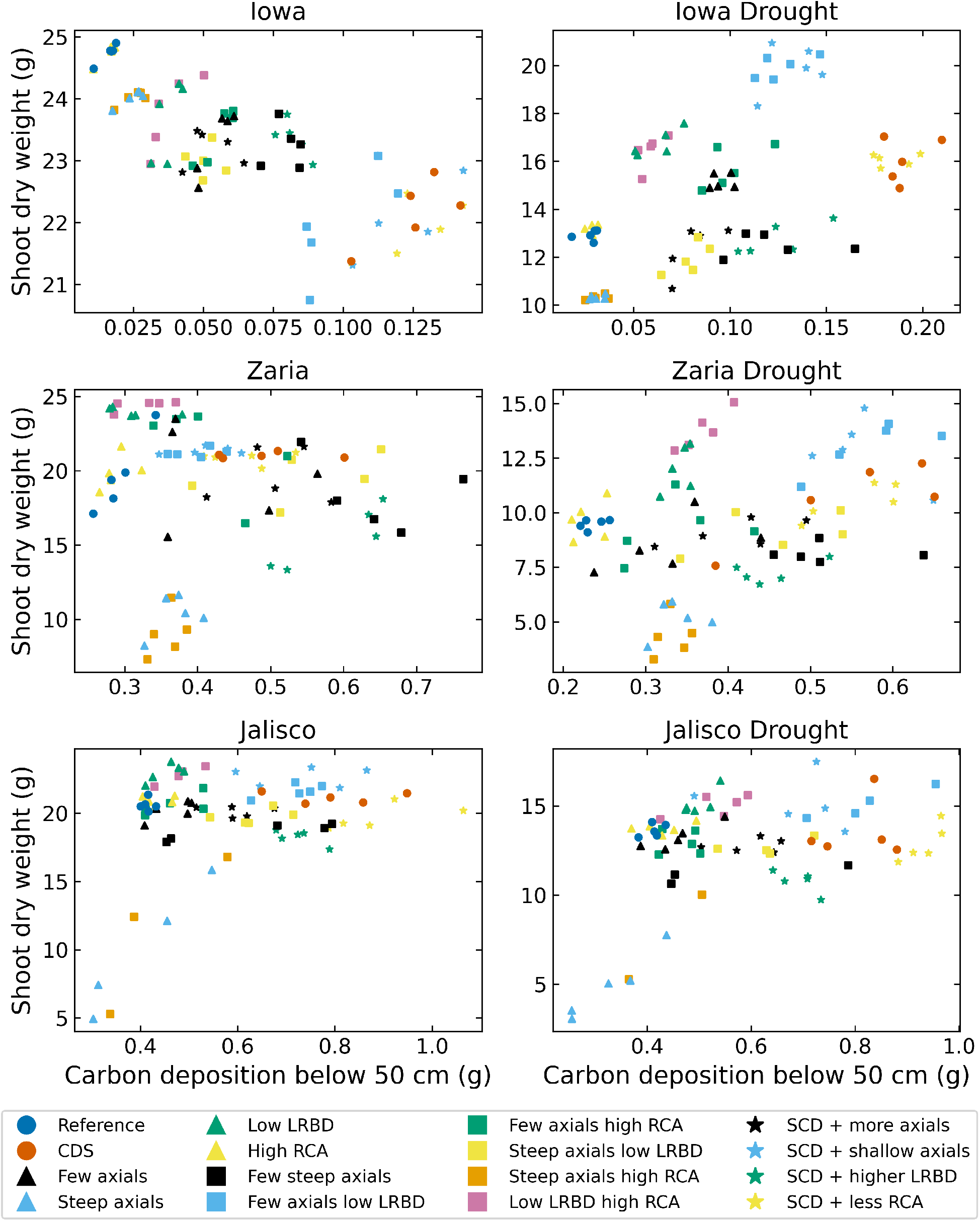
Carbon depositions below 50 cm deep in the soil versus shoot dry weight of all the maize phenotypes considered after 42 days in the 6 different environments. LRBD means lateral root branching density, RCA means root cortical aerenchyma and SCD refers to the steep, cheap, deep phenotype.

Plants with the SCD phenotype had less water capture in the upper 40 cm of soil and greater water capture in deeper soil strata (Figure 10). Plants with the SCD phenotype also acquired water from deeper in the soil. The difference in water capture from the subsoil was greater in drought environments than in rainfed environments. In some environments, negative total water uptake in the topsoil highlights the occurrence of hydraulic lift.

**Figure 10.**
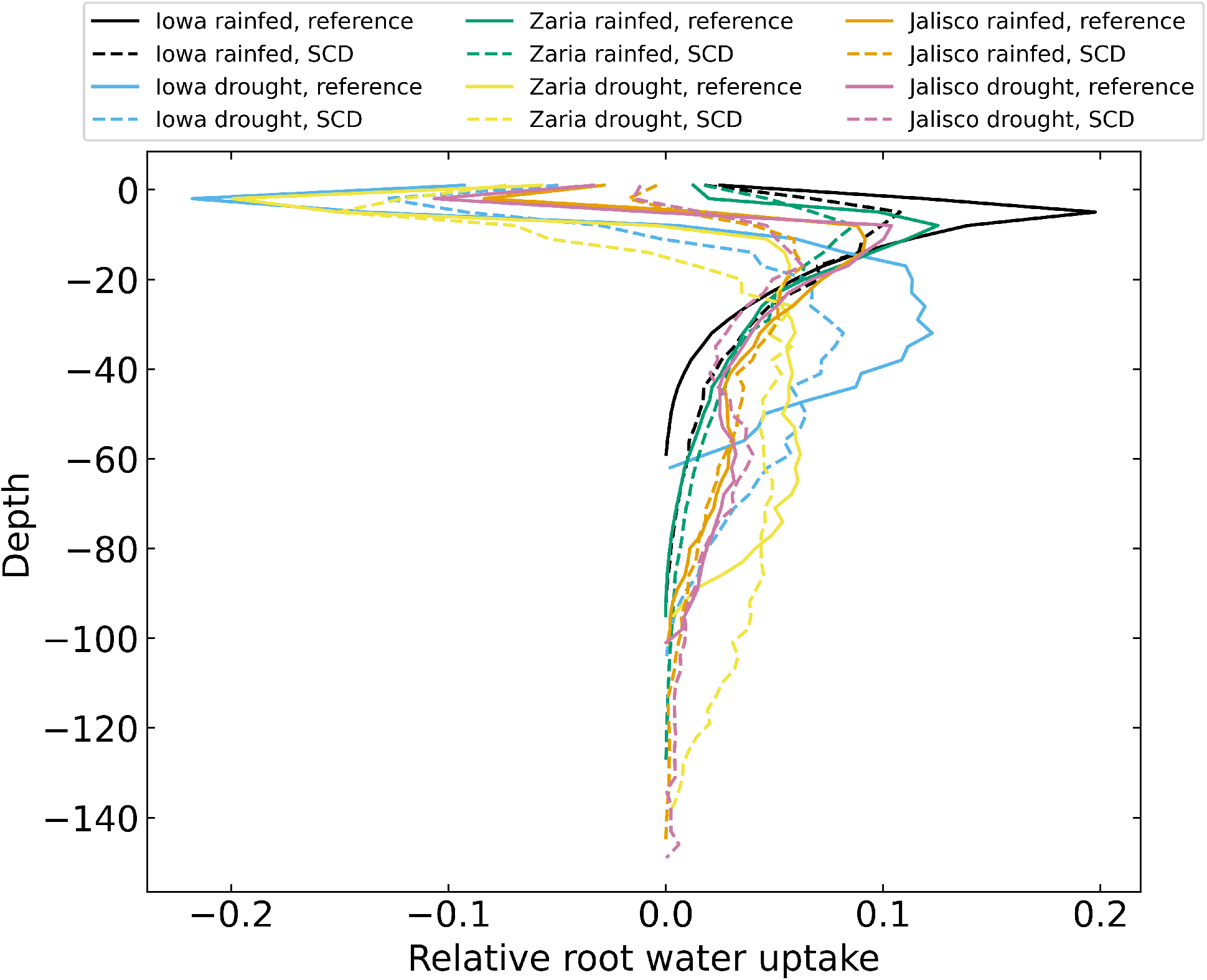
Relative root water uptake after 42 days for 12 individual simulations, averaged over 3 cm layers. There is negative uptake in the upper soil layers because of hydraulic lift.

We observed substantial synergism (i.e., greater benefit than expected from simply additive interaction) among root phene states for plant performance in all environments except rainfed conditions in Iowa (Figure 11). The magnitude of interactive effect was substantial, ranging from antagonism of -36% (ACD, or SCD plus shallow axial roots in rainfed Zaria) to synergism of +226% (ABD, or SCD plus greater lateral root branching density in Jalisco under drought), averaging +30% over the additive effects. In Zaria and Jalisco, 6 integrated phenotypes showed strong synergism in multiple environments, of which all 6 include steeper axial root growth angles, 4 include fewer axial roots and 4 include reduced lateral root branching density. In Iowa under drought, two integrated phenotypes displayed synergism, both containing the combination of steeper axial root growth angles and increased aerenchyma formation. Synergism was greater under drought than under rainfed environments, averaging +39% under drought compared with +24% under rainfed conditions in Zaria, and +74% under drought compared with +42% under rainfed conditions in Jalisco, all relative to the additive interactions (Figure 11). Synergism was smaller and less common in the Iowa environment.

**Figure 11.**
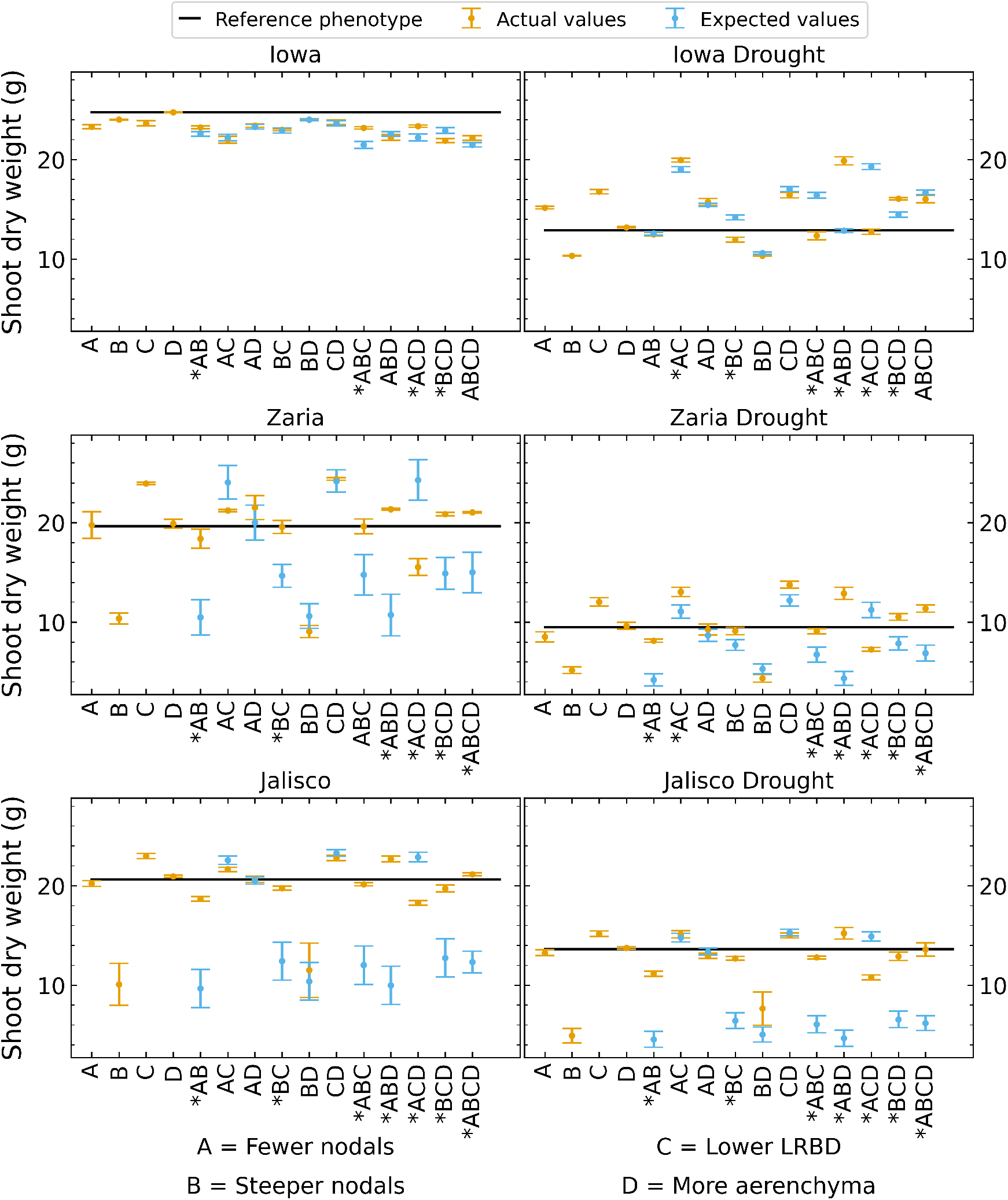
Shoot dry weight of phenotypes as expected by adding the changes resulting from altering individual phenes compared to the actual shoot dry weights of these phenotypes. The black line indicates the shoot dry weight of the reference phenotype. The error bars indicate standard error. Cases where the expected and actual shoot dry weight are significantly different are marked by an asterisk on the x axis label.

## 4 DISCUSSION

The implementation of established biophysical-biochemical models to mechanistically capture various shoot processes in the functional-structural plant/soil model *OpenSimRoot* has led to the development of the most feature-rich root/soil modeling platform, *OpenSimRoot_v2*, available in the literature to date, to our knowledge,. Using the *OpenSimRoot_v2* model, we evaluated the utility of SCD root phenotypes under rainfed and drought conditions over three distinct pedo-climatic environments to evaluate the relationship of root phenotypes with maize growth and soil carbon deposition under drought. Our results show that parsimonious root phenotypes with fewer axial and lateral roots, improve shoot dry weight significantly in terminal drought conditions, as hypothesised Lynch (2013); Passioura (1983). We observe that the SCD phenotype spends less carbon on the root system without negatively affecting water capture. This results in greater carbon use efficiency (water acquired per unit of carbon invested in roots) that in turn leads to greater shoot biomass. The increase in root length at depth allows plants with the SCD phenotype to access deeper water, and distribute water capture over more soil strata, so roots encounter softer, wetter soil, and increase deep soil carbon deposition.

Reduced axial root number has been suggested as a target for breeding more drought-resistant plants Lynch (2018). Maize plants with fewer crown roots had greater shoot biomass under drought in mesocosms and the field Gao and Lynch (2016). Reducing axial root number increased shoot dry weight in the Iowa drought environment but it slightly reduced or had little effect on shoot dry weight in the other environments (Figure 2).Reducing axial root number decreased root carbon costs and increased root length and relative root length below 50 cm (Figures 3 5). This reduced root carbon cost and a deeper root system were not associated with an increase in shoot dry weight.

Having steeper axial roots by itself was detrimental in all 6 of the simulated environments (Figure 2).This result highlights the importance of capturing water available in the topsoil, even in drought environments where more water is found in the subsoil. Shallow root systems are subject to greater interplant competition for topsoil resources like phosphorus Rubio et al. (2001); Lynch and Brown (2001) while steeper root systems experience greater intraplant competition for mobile resources like nitrogen and water Nord et al. (2011); Trachsel et al. (2013); Ajmera et al. (2022). Steeper axial root angles are associated with deeper root systems Oyanagi et al. (1993); Uga et al. (2015), but the benefit of steeper axial roots depends on environmental factors such as the availability of subsoil water and the amount of in-season precipitation Manschadi et al. (2008). Root systems with both greater root length density in intermediate and deep soil layers and deeper maximum depth are candidates for increased subsoil water uptake and thus drought resistance Hund et al. (2009); Wasson et al. (2012). Making the axial root angles of the reference phenotype steeper increased root length below 50 cm, but markedly less than doing this in combination with reducing axial root number and lateral branching density, which highlights the importance of interactions between root phenotypes and the environment in determining drought resistance Ludlow and Muchow (1990); Richards et al. (2002). While increased root length in the subsoil was beneficial in the terminal drought scenario, large and shallow root systems were beneficial in some simulated environments. Capturing water in the topsoil, which is of course where precipitation enters the soil, will be even more important for crops that have to compete with weeds and intercrops or cover crops for limited water resources.

Reducing lateral root branching density increased shoot dry weight in all environments except the rainfed Iowa environment (Figure 2). Since competition among neighboring lateral roots for water is significant, reducing lateral branching density reduces root system carbon costs much more than it reduces water capture. In fact, the reduction in lateral root carbon costs due to reduced lateral branching density increased photosynthate available for axial roots, which were then able to grow deeper, increasing root depth (Figure 5). These results are consistent with the report that reduced lateral root branching density was associated with deeper rooting, better capture of subsoil water, and better plant water status, growth and yield in maize under drought stress in two field environments and in greenhouse mesocosms Zhan et al. (2015). Similarly, maize lines with reduced lateral root branching density had deeper rooting, greater plant N content, photosynthesis, growth, and yield under low N stress in two field environments and in greenhouse mesocosms Zhan and Lynch (2015).

Increasing aerenchyma formation had little effect on shoot dry weight in all environments (Figure 2). Likewise, for most intermediate phenotypes to which increased aerenchyma formation was added, shoot dry weight did not change much. This contrasts with field studies where increased aerenchyma formation was associated with substantially greater maize yield under drought in the field Chimungu et al. (2015); Zhu et al. (2010). This discrepancy is perhaps due to the narrow range of increase in aerenchyma formation (i.e., from 20% to 40%) considered in this study compared to the field experiments wherein aerenchyma formation was observed to vary from 0 to 50% in maize under drought with significant stress and environmental plasticity Schneider et al. (2020). Aerenchyma influences plant performance by reducing root respiration and remobilizing tissue nutrients, which are explicitly simulated in the model. Since nutrient constraints on growth are not included in this drought study, the benefit of aerenchyma formation is less than that observed in response to suboptimal nutrient availability Postma and Lynch (2011).

Our observation that phene synergism is an important component of the fitness of integrated root phenotypes supports previous reports of substantial phene synergism for soil resource capture *in silico* and *in planta*. For example, increased root hair length, root hair density, root hair proximity to the root tip, and number of trichoblast files increased P acquisition in Arabidopsis roots by 371% more than their additive effects *in silico* Ma et al. (2001). In simulated maize roots, Root cortical aerenchyma increased P acquisition 2.9 times more in phenotypes with increased lateral branching density Postma and Lynch (2011). Multiple synergies are evident in the interactions of axial root phenotypes of common bean for the acquisition of N and P in silico *Rangarajan et al. (2018). In planta*, metaxylem anatomy and root depth are synergistic for drought adaptation in contrasting *Phaseolus* species Strock et al. (2021), and shallow root growth angles and long root hairs are synergistic for P capture in common bean Miguel et al. (2015). Here, we attribute synergism between fewer axial roots and steeper axial root growth angles to reduced intraplant competition for water capture. Steeper root growth angles increase subsoil exploration but also position root axes closer together, thereby increasing competition among root axes for soil resources, especially mobile resources such as water Dathe et al. (2016). Two synergistic integrated phenotypes combine steeper axial root growth angles with reduced lateral root branching density, which was synergistic in 4 of the 6 environments. Reduced lateral root branching, as with reduced axial root number, would reduce competition among roots for soil water. In this context we note that slight water stress occurs under our rainfed Zaria and Jalisco scenarios (Figure 2), meaning that root phenotypes that improve water capture should improve plant growth under both rainfed and drought environments. We interpret the smaller degree of synergism observed in the Iowa environments to the significantly greater bulk density of the Iowa soil, which inhibits soil penetration by small diameter roots, reducing intraplant competition for water capture.

While shoot dry weight under drought correlates very strongly with water uptake, we also found a strong relationship between carbon use efficiency (water uptake per unit of carbon invested in roots) and shoot dry weight. A similar metric, root system efficiency (transpiration per unit leaf area per unit of root mass) was proposed as breeding target for drought resistant maize van Oosterom et al. (2016). By reducing the amount of carbon used for growing and maintaining the root system, a plant frees up carbon which can be used to grow a bigger shoot or for meeting metabolic needs during drought stress Bolaños et al. (1993); L ynch (2018); Passioura (1983). If the carbon required to grow extra roots do not result in enough additional water uptake to produce more carbon, the plant is effectively moving biomass underground, reducing its yield potential. Three of the 4 phene states included in the SCD phenotype reduce root system carbon costs: Fewer axial roots, reduced lateral branching density and greater root cortical aerenchyma formation. Noting that reducing the axial root number and lateral branching density increase carbon use efficiency more than increased aerenchyma formation, we conclude that competition among roots is significant in the reference phenotype. Besides increasing root cortical aerenchyma formation, other anatomical phenotypes, such as reduced cortical cell file number and increased cortical cell size reduce root respiratory demands, potentially increasing carbon use efficiency even more if integrated with the SCD phenotype Lynch (2015).

Besides improving plant development under drought to mitigate the impact of climate change on yield, it has been suggested that improved carbon storage in the soil by plant roots would sequester atmospheric CO_2_ Dondini et al. (2009); Kell (2011, 2012); Lorenz and Lal (2005). Due to the reduced presence of oxygen and biological activity, carbon recalcitrance increases with soil depth Lorenz and Lal (2005). While *OpenSimRoot* does not yet simulate the carbon cycle in the soil, we can calculate the carbon deposited below 50 cm in the soil in the form of roots and exudates. In all environments, the SCD phenotype significantly increased carbon deposition in deep soil. Increasing the carbon deposition by at least 0.3 gram per plant, as we observed in Zaria and Jalisco, at the planting density of 8 plants per m^2^ amounts to deposition of 24 kg of carbon, or removal of approximately 88 kg of CO_2_ per hectare. This increase of 0.3-gram carbon per plant deposited does not take into account respiration and is observed after 42 days of growth, so the actual amount after harvest (80 to 160 days) will be greater. Our simulations suggest that the SCD ideotype could increase shoot dry weight as well as carbon deposition in deep soil.

Our results highlight the importance of soil-plant interactions when modeling crop development since we see major differences among the three soils. Soil compaction can negatively impact root growth and plant development Taylor and Brar (1991); Unger and Kaspar (1994). In the Iowa soil, the soil bulk density below 45 cm is greater than in the other soils, which leads to large differences in root length in deep soil strata, as compared to the other two soils. Where the reference phenotype has at least 1000 cm and up to 3000 cm of root length below 50 cm in Zaria and Jalisco, in Iowa, it was less than 100 cm. Soil compaction caused by long term use of heavy agricultural machinery, as occurs in the Iowa environment, makes it more difficult for axial roots to reach deeper soil layers, and leads to a reduction of lateral root growth in the compaction layer, resulting in a more shallow root system. This is consistent with the results of a field study where 94% of total maize root length was located in the upper 60 cm of soil, with 84% in the top 30 cm, and water uptake mainly occurred in the upper 75 cm of soil Laboski et al. (1998). Rather than the maximum rooting depth, the depth of the most densely rooted soil layer was important for maize plants to cope with drought Yu et al. (2007). Irrespective of the presence of a compaction layer, the SCD phenotype increases subsoil root length significantly (rather than just the maximum rooting depth), explaining the increased drought resistance.

The soil affects the way roots grow and in turn, roots affect soil properties. Plants growing in compacted soil will have a more shallow root system and the water they take up from the topsoil increases soil penetration resistance so that reaching deeper soil layers where water might still be available becomes more difficult Colombi et al. (2018). Our results suggest that the SCD ideotype leads to greater water content near the roots than the reference phenotype at most depths, resulting in reduced soil penetration resistance. In drought environments, the reference phenotype has greater water content around the deepest roots than the SCD phenotype had around the same depth, but the SCD phenotype had similar water content in deeper soil strata (the SCD phenotype had deeper roots). This makes sense because root length density will be very low at the deepest point of a root system, so the plant will not take up much water there. Because the SCD phenotype is deeper than the reference phenotype, it has access to deeper soil water and water capture has greater vertical distribution. In this way, the SCD ideotype can help plants avoid the feedback loop (shallow root systems take up more water from the topsoil, which increases topsoil penetration resistance, making it harder for roots to reach the subsoil) described in Colombi et al. (2018). Steeper axial roots mean that a root system is less wide but at commercial planting densities a wide root system will lead to increased competition with neighboring plants and is unlikely to increase water capture at the stand scale.

If plants with the SCD root phenotype perform better under drought without any tradeoffs one would expect maize plants to already possess this phenotype. That substantial variation exists for root phenotypes in maize suggests that a single root phenotype may not be optimal in all conditions. If water and nitrate are readily available then the benefit of the SCD ideotype for water and N acquisition is irrelevant, but the reduced carbon demands remain beneficial. Previous studies have shown that in soils with low phosphorus availability, high root length density in the topsoil is beneficial Lynch and Brown (2001); Postma et al. (2014); Rubio et al. (2003). While nitrate is mobile in the soil, if precipitation rates are low then nitrate will remain concentrated in the topsoil and shallow root systems perform better Dathe et al. (2016). It has been suggested that plants with an SCD phenotype may also be more susceptible to root loss caused by diseases and herbivory, because each root represents a larger fraction of the total root system Schäfer et al. (2022a).

The results presented here should be understood to be obtained with a number of limitations. As in any modelling study, many processes have been simplified or omitted. *OpenSimRoot* does not simulate phytohormones. While care was taken to include factors relevant to modeling photosynthesis under drought such as canopy temperature, canopy self-shading, light scattering, and stomatal conductance in response to plant water status, the model is inevitably a simplification of reality. We assumed the stomata can close and open instantly and the canopy is always at the equilibrium temperature. Due to computational constraints, we are simulating at finite temporal and spatial resolutions: the simulations run at a timestep of 0.1 days, which means we simulate the plant at different times of day (morning, noon, afternoon, etc.) while keeping computational costs manageable. The soil has a resolution of 1 cm, which allows for some differentiation between the resource capture of different roots but means the soil water gradient around individual roots cannot be accurately simulated. A limitation to the current model is that while the water flowing out of the roots during a hydraulic lift is simulated, this water is not added back to the soil, which means the topsoil has a less water content than it should. The total outflow is only a few percent compared to the total water content in the soil or the total change in soil water over the 42 simulated days so this will have a minor effect on results.

Perhaps the greatest limitation is that plants are only simulated for 42 days instead of until harvest. This is because the exponential increase in root segments over time means that computational costs of every simulated day increase as the root system grows. Simulating a maize plant for 42 days requires several days of computational time in some cases. While shoot dry weight correlates with yield Hay (1995); Huetsch and Schubert (2017), the relationship between them is nonlinear, especially under drought stress, and there is no guarantee that the phenotypes with the greatest shoot dry weight at 42 days will also have the greatest shoot dry weight at harvest. Therefore, our results are relevant to vegetative growth under drought rather than yield under drought. An important factor under terminal drought is the banking of soil water to sustain reproductive growth Zhang et al. (2019), a process not simulated here. Likewise, for carbon sequestration, the final amount of carbon deposited in the soil is relevant and it is possible a plant starts off slowly but deposits more carbon overall. However, since we see a strong correlation between shoot dry weight and carbon use efficiency under drought, the qualitative conclusions drawn from our results should be valid even when older plants are considered.

In summary, *OpenSimRoot_v2* is capable of simulating maize growth under water deficit stress, encompassing root exploration of drying, hardening soil, water acquisition, and photosynthetic responses to water availability. Implementing this model, we present *in silico* evidence that more parsimonious root phenotypes, as epitomized by the SCD ideotype, that optimize the metabolic costs of soil exploration and water acquisition are capable of improving vegetative growth and carbon deposition in deep soil under drought in distinct maize production environments.

## AUTHOR CONTRIBUTIONS

ES, IA and JL contributed to conception and design of the study. ES and IA selected, implemented and tested the models, ran the simulations and analyzed the data. EF, LB and MO supervized model implementation and helped resolve modelling issues. ES wrote the first draft of the manuscript. ES, IA and JL wrote sections of the manuscript. All authors contributed to manuscript revision, read and approved the submitted version.

## ACKNOWLEDGMENTS

The authors are grateful for access to the University of Nottingham High Performance Computing Facility. This study also used Extreme Science and Engineering Discovery Environment resource - Bridges at Pittsburgh Supercomputing Centre through allocation BCS-200008. The authors acknowledge Johannes A.

Postma, Christian Kuppe (Forschungszentrum Juülich), and Nathan Mellor (University of Nottingham, UK) for their support during the initial development of the model.

## SUPPLEMENTAL DATA

The following materials are available in the online version of this article.

Supplementary Information 1: Link to reference input file and the *OpenSimRoot_v2* executable employed in this study.

## DATA AVAILABILITY STATEMENT

Data will be available from the corresponding author on request

